# eIF3d drives cholesterol biosynthesis required for KSHV lytic replication

**DOI:** 10.1101/356162

**Authors:** Eric S. Pringle, Carolyn-Ann Robinson, Brett A. Duguay, Christopher S. Hughes, Nicolas Crapoulet, Andrea L-A. Monjo, Alexa Wilson, Katrina Bouzanis, Andrew M. Leidal, Stephen M. Lewis, Daniel Gaston, James Uniacke, Craig McCormick

## Abstract

Herpesvirus genomes are decoded by host RNA polymerase II, generating viral messenger ribonucleic acids (mRNAs) that are post-transcriptionally modified and exported to the cytoplasm. These viral mRNAs have 5□-m^7^GTP caps and poly(A) tails that enable assembly of host translation initiation factors (TIFs) required for viral protein synthesis. Here we show that maximal production of viral proteins during lytic replication requires both the canonical host TIF eukaryotic initiation factor 4F (eIF4F) and the alternative host TIF eIF3d. Despite this, production of infectious virions is largely dependent on eIF3d, with eIF4F playing a minor role. Investigating the effects of *eIF3d* silencing on the host proteome revealed that it is required for accumulation of proteins involved in cholesterol metabolism, as well as the accumulation of intracellular cholesterol during KSHV lytic replication. In KSHV infected and uninfected cells alike, eIF3d supports efficient translation of cholesterol biosynthesis enzymes, including squalene epoxidase (SQLE), whose function is required to support productive lytic replication during cholesterol scarcity. These findings position eIF3d as a critical TIF required to support translation of essential host mRNAs required for KSHV replication.

**AUTHOR SUMMARY:** Animal cells tightly regulate uptake, synthesis, storage, and export of cholesterol to ensure optimal membrane fluidity and permeability. Many enveloped viruses require cholesterol to support membrane fusion events during cell entry and cell surface budding events during assembly and egress. Herpesviruses like Kaposi’s sarcoma-associated virus (KSHV) require cholesterol to support entry, but its role in post-entry events, including budding and fusion at internal membranes, remains incompletely understood. Here, through our studies of KSHV protein synthesis, we uncovered a new mechanism of cellular control of cholesterol metabolism involving the host translation initiation factor (TIF) eIF3d. We demonstrated that eIF3d was required to support efficient translation of host mRNAs encoding key proteins required for cholesterol biosynthesis and uptake. Using a lytic reactivation model that bypasses the entry step, we observed that productive KSHV lytic replication is accompanied by eIF3d-dependent increases in intracellular cholesterol. Preventing cholesterol synthesis and uptake, or silencing *eIF3d*, strongly inhibited production of infectious virions. These studies shed new light on the importance of intracellular cholesterol to support KSHV replication, and provide motivation for future studies of how cholesterol influences discrete post-entry steps of KSHV infection.

## INTRODUCTION

All viruses use host translation machinery to synthesize viral proteins. DNA viruses, such as herpesviruses, replicate in the nucleus where they can access the full complement of host transcription machinery ^1–3^. Thus, herpesvirus mRNAs are transcribed by RNA polymerase II (Pol II) and processed by enzymes that add 5□ m^7^GTP caps and 3□ poly-adenylate (polyA) tails, features that promote mRNA stability and provide access to the same cap-dependent translation initiation factors (TIF) employed by cellular mRNAs ^2,4^. In typical cell culture experiments (normoxia, rich culture media, hard plastic substrate), approximately two-thirds of global protein synthesis requires eukaryotic initiation factor 4F (eIF4F) ^5^, a protein complex comprising the eIF4E m^7^GTP cap-binding protein, the eIF4G scaffolding protein, and the eIF4A RNA helicase ^6–9^. After the cap is bound by eIF4E, eIF4G recruits eIF3 and the small ribosomal subunit to initiate scanning for the start codon. eIF4A with its cofactors eIF4B and eIF4H reduce secondary structure in the 5□ untranslated region (UTR) to facilitate this process ^7,9^.

In response to nutrients and growth signals, mechanistic target of rapamycin complex 1 (mTORC1) regulates translation initiation by promoting eIF4F assembly. When unphosphorylated, repressive 4E-BP1 proteins bind to the eIF4E cap-binding protein and prevent recruitment the eIF4G scaffolding protein required for eIF4F assembly. mTORC1-mediated phosphorylation of 4E-BP1 liberates eIF4E and enables eIF4F assembly and subsequent recruitment of the eIF3 complex and the small ribosomal subunit ^3,8,8,10^. mTORC1 inhibition causes widespread decreases in protein synthesis, but transcripts bearing 5□ terminal oligopyrimidine (TOP) sequences ^6^ or pyrimidine-rich translational element (PRTE) sequences ^11^ at their 5□ ends are especially sensitive to mTORC1 inhibition. When eIF4F assembly is inhibited in response to nutrient scarcity or treatment with mTORC1 inhibitors, residual translation can be supported by eIF3d, a subunit of eIF3 that contains a cap-binding domain and can direct cap-dependent translation ^5,12–14^. The cap-binding domain of eIF3d is normally blocked by phosphorylation by casein kinase 2 (CK2) in fed cells, but this phosphorylation is lost when glucose is scarce, indicating that access to eIF3d is metabolically regulated ^13^.

Herpesviruses from all subfamilies (alpha-, beta-, and gammaherpesviruses) have been shown to activate mTORC1 during lytic replication ^15–18^, but paradoxically, virion production is only modestly reduced by mTORC1 inhibition ^17,19,20^. We and others have shown that eIF4F assembles and remains sensitive to the mTORC1 active-site inhibitor Torin throughout the Kaposi’s sarcoma-associated herpesvirus (KSHV) lytic replication cycle ^15,17,21^. These findings suggest that translation of KSHV mRNAs can be supported by alternative non-eIF4F host TIFs during lytic replication. Here, to better understand the determinants of KSHV protein synthesis in the context of lytic replication, we used polysome profiling to measure translational efficiency (TE) of viral mRNAs and proteomics to measure accumulation of viral proteins. We discovered that eIF4F and eIF3d both contribute to the accumulation of viral proteins during KSHV lytic replication. However, eIF3d depletion greatly diminishes release of infectious virions even when eIF4F is available to compensate for lost translation initiation capacity. We observed that *eIF3d* silencing reduced the accumulation of host proteins that regulate cholesterol metabolism and prevented the accumulation of intracellular cholesterol during KSHV lytic replication. In KSHV infected and uninfected cells alike, eIF3d supported efficient translation of cholesterol biosynthesis enzymes including squalene epoxidase (SQLE), whose function was required to support productive lytic replication during cholesterol scarcity. These findings position eIF3d as a critical TIF required to successfully complete the viral replication cycle.

## RESULTS

### eIF4F is dispensable for translation of KSHV lytic mRNAs

We and others previously reported that eIF4F assembles during KSHV latency and lytic replication and remains under strict regulation by mTORC1 and 4E-BP1 ^15,17^. Here, we used polysome profiling to measure how eIF4F disruption affected translational efficiency (TE) of viral mRNAs. Physiologic mTORC1 activity is regulated by nutrient abundance, but mTORC1 is readily inhibited by several drugs, including the potent active site inhibitor Torin ^21^. We observed that Torin treatment of uninfected iSLK cells for 2 h prior to harvest caused dephosphorylation of ribosomal protein S6 and 4E-BP1 as expected, which was likewise evident in latently KSHV-infected iSLK.219 cells, or cells which had been treated with doxycycline (dox) for 48 h to activate the lytic cycle (**Fig. 1A**). In these experiments, we reactivated iSLK.219 cells with dox only; we omitted the HDAC inhibitor sodium butyrate that is commonly used to promote lytic reactivation to limit potentially confounding effects of epigenetic regulation of transcription by histone acetylation. While we observe incomplete reactivation under these conditions, there is still significant cytopathic effect (CPE) at 48 h and 72 h and there is near complete clearing of the monolayer by 96 h, as reported previously ^22^. mTORC1 regulation of canonical target proteins S6 and 4E-BP1 remains largely intact during latent and lytic phases of KSHV replication in this model. As expected, we observed that treatment of uninfected iSLK cells with Torin decreased heavy polysomes and increased monosome fractions (**Fig. 1B, top panel**) consistent with diminished bulk protein synthesis, as had been observed in studies of other cell types ^6,20^. Latently infected iSLK.219 cells treated with Torin displayed a similar shift of mRNA and ribosomes into monosomal fractions (**Fig. 1B, middle panel**). By 48 h after reactivation from latency, heavy polysomes were reduced (**Fig. 1B, bottom panel**), likely due to host shutoff ^23^; Torin treatment further depleted the heavy polysome fractions in these cells, although the effects were modest, possibly due to the pre-existing low translation environment in these cells.

**Fig 1.**
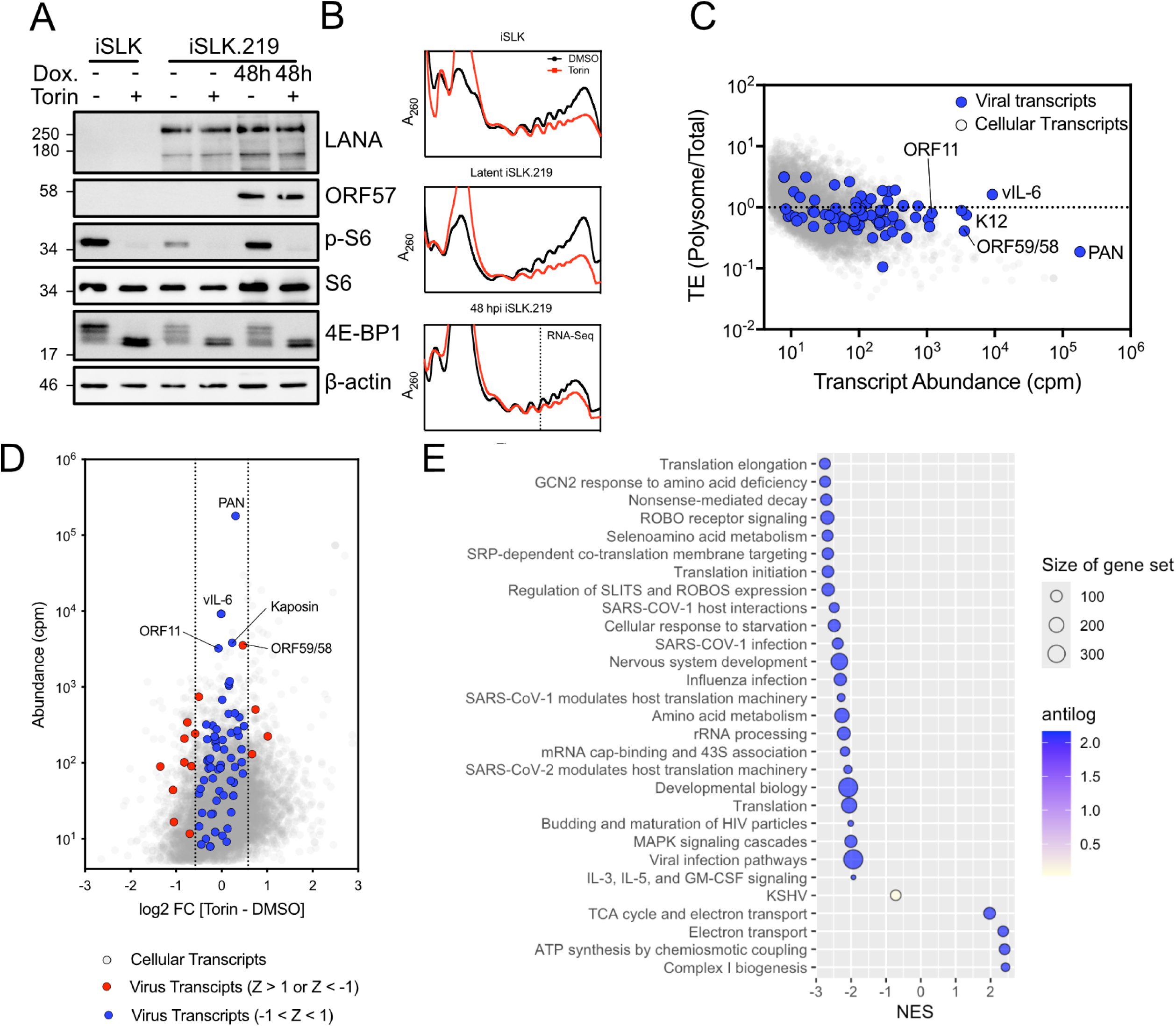
eIF4F disassembly does not affect translational efficiencies of viral mRNA. **(A)** Western blots of whole cell lysates of uninfected iSLK, latently infected iSLK.219 and 48 h post-doxycycline (Dox.) treatment (hours post-induction (hpi)). iSLK.219 cells treated with 250 nM Torin or DMSO vehicle control for 2 h prior to harvest. **(B)** Polysome profiles of uninfected, latent, or 48 hpi iSLK.219 treated with either Torin or DMSO for 2 h prior to harvest. **(C)** mRNA from translating ribosomes (of DMSO treated cells) was sequenced and aligned to both the human and KSHV genomes. Viral transcripts are depicted in blue on top of the grey background of cellular genes. **(D)** The change in translational efficiency (Δ TE) of viral transcripts is depicted in blue or red on a grey background of cellular genes. Viral transcripts beyond one SD of the mean are red, viral transcripts within one standard deviation (SD) are depicted in blue. Vertical lines represent a 1.5-fold change in transcript TE. **(E)** Gene set enrichment analysis of transcripts TE < -1.5 Z or TE > 1.5 Z.

To measure the effects of mTORC1 inhibition on the TE of viral and host mRNAs during the KSHV lytic cycle, we isolated RNA from polysome fractions from Torin- and vehicle-treated cells at 48 h post-reactivation (**Fig. 1B, bottom panel**) and processed it for RNA sequencing. We assessed TE by simple division of the number of reads isolated from the polysomes compared to the reads found in the total RNA fraction (# of reads polysome / # of reads total). The abundance of KSHV transcripts in the total RNA fraction differed by as much as 100,000-fold, with ten-fold more of the non-coding PAN RNA than the next most abundant RNA **(Fig. 1C, S1 Table)** ^2,24^. Despite these differences, the TE of most viral mRNAs was similar to most cellular mRNAs. Consistent with the literature, we observed alterations in TE of cellular mRNAs in the presence of Torin, with populations of mRNAs displaying increased or decreased TE **(Fig. 1D, S2 Table)**. The ΔTE of most viral mRNAs (∼83%) was not inhibited or enhanced by a conservative Z-score of 1 (Z-score within 1 SD of the mean in blue, Z-score > 1 SD of the mean in red). We analysed transcripts with a greater than 1.5-fold change in TE using Gene Set Enrichment Analysis (GSEA) to determine how the translation of cellular mRNAs is altered by eIF4F depletion during lytic replication ^25^ **(Fig. 1E)**. Gene sets involved in translation elongation and initiation featured prominently in transcripts with reduced TE, consistent with previous studies ^6,11^, which suggests that TOP-containing transcripts are regulated by mTORC1 as expected during KSHV lytic replication. Notably, the TE of a gene set of viral mRNAs was not significantly altered by Torin treatment. Together, these data suggest that even though normal regulation of translation by the mTORC1/4E-BP/eIF4F axis is intact during lytic replication, viral mRNAs continue to be efficiently translated when eIF4F is depleted.

### eIF3d is required for production of infectious KSHV virions

When eIF4F assembly is prevented by mTORC1 inhibitors, residual translation can be supported by eIF3d ^5,12–14^. To assess the importance of eIF3d in supporting KSHV lytic replication, we used short interfering RNAs (siRNAs) to silence *eIF3d* **(Fig. 2A)** and measured production of infectious virions at 96 h post-reactivation, in the presence or absence of Torin. We previously demonstrated that mTORC1 activity is required to support the earliest stages of lytic replication, so Torin was added at 24 h post-reactivation to coincide with the transition to mTORC1-independent viral replication ^17^. Torin treatment from 24 h to 96 h only elicited a modest reduction in production of infectious virions (**Fig. 2B**), as we have observed previously ^17^. By contrast, when *eIF3d* was silenced, there was a ∼4 log_10_ reduction in titer **(Fig. 2B)** with very little change in the abundance of viral mRNA **(Fig. 2C),** suggesting a defect in viral translation even when eIF4F is available. We corroborated this finding with independent short hairpin RNAs (shRNAs) encoded by lentiviral constructs, which also effectively silenced *eIF3d* (**Fig. 2D**) and inhibited production of infectious KSHV virions (**Fig. 2E**).

**Fig. 2.**
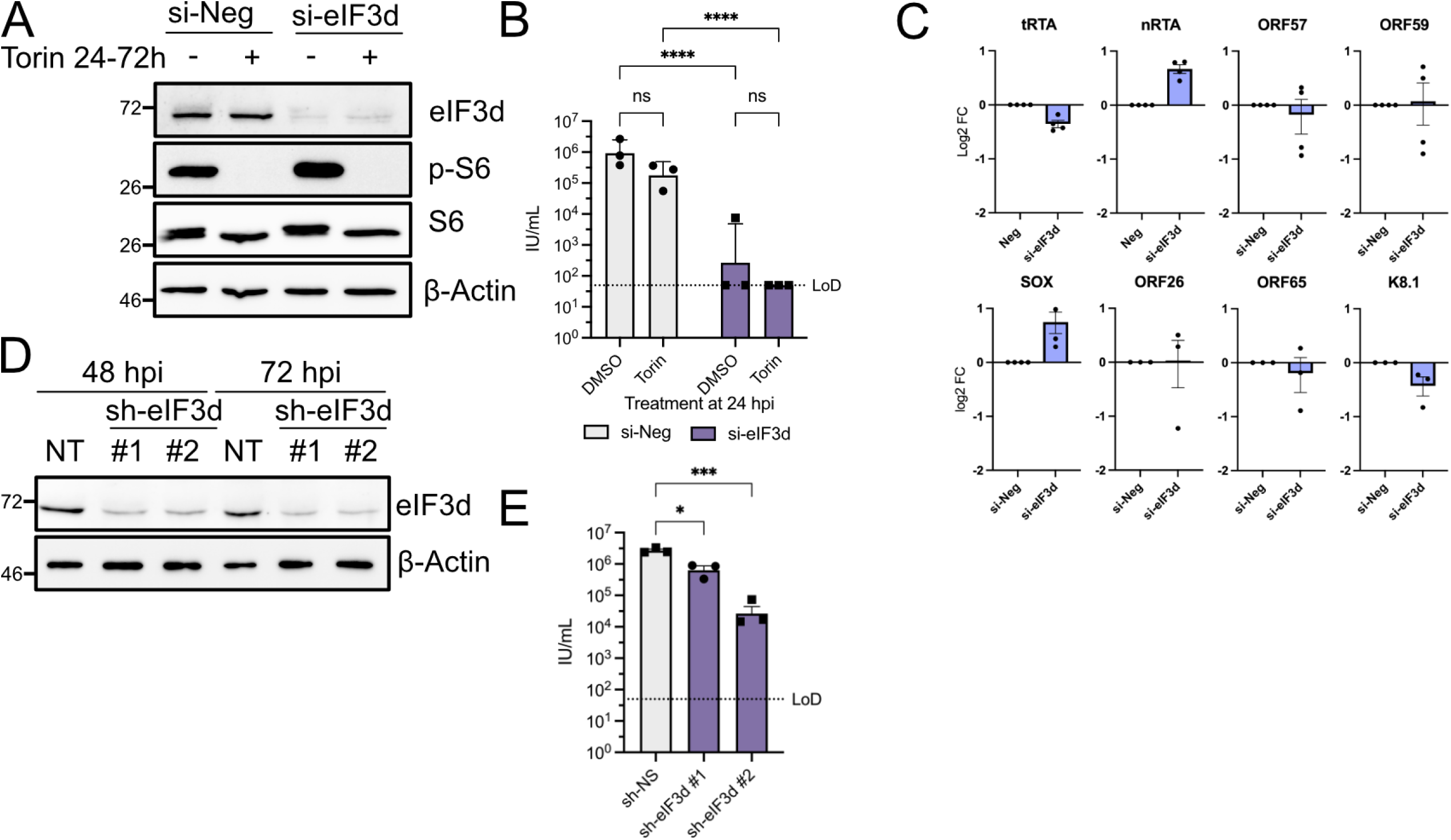
eIF3d is required for accumulation of KSHV lytic protein products. **(A)** Western blot of whole cell lysates of 72 h post-doxycycline (Dox.) treatment (hpi) previously transfected with siRNA targeting eIF3d (si-eIF3d) or a negative control siRNA (si-Neg). Cells were treated with 250 nM Torin or DMSO vehicle control at 24 hpi **(B)** Viral titer at 96 h post-reactivation from iSLK.219 cells treated as in **(A)** (n=3; mean ± standard error of the mean (SEM); statistical significance was determined by two-way ANOVA). **(C)** RNA was harvested at 72h post-reactivation in siRNA-treated cells for RT-qPCR using primers targeting viral gene products as indicated (n=4; mean ± SEM; statistical significance was determined by two-way ANOVA). **(D)** Western blot of whole cell lysates of latently infected iSLK.219 transduced with lentivirus encoding shRNA targeting eIF3d (sh-eIF3d #1 and #2) or a non-targeting negative control shRNA (NT) in cells at 48 or 72 hpi **(E)** Viral titer at 96 h post-reactivation from iSLK.219 cells treated as in **(D)** (n=3; mean ± SEM; statistical significance was determined by two-way ANOVA). Abbreviations: nRTA, native RTA; ns, not significant; tRTA, transgenic RTA.

Using the same experimental conditions (*eIF3d* silencing +/- Torin treatment beginning at 24 h post-reactivation), we assessed the accumulation of viral proteins by label-free proteomics, harvesting total protein at 72 h post-reactivation, a time point that coincides with abundant late protein accumulation, following genome replication and preceding bulk virion release ^17^. We observed that Torin treatment had no effect on eIF3d abundance (**Fig 3A, green circle, S3 Table**), whereas the siRNA targeting *eIF3d* potently and significantly reduced total eIF3d protein (**Fig. 3B, green circle, S4 Table**), validating our RNA silencing approach. Both Torin treatment and *eIF3d* silencing reduced the accumulation of most viral proteins at 72 h (depicted in red on the volcano plots; **Figs. 3A, 3B**). However, we noted differences in the specific proteins that were downregulated by inhibiting each translation initiation system (**Fig. 3C, S5 Table**), suggesting slightly different effects on the translation of these mRNAs. Interestingly, combining both Torin treatment and eIF3d knockdown resulted in an even greater suppression of viral protein accumulation (**Fig. 3D, S6 Table**). To determine the effect of *eIF3d* silencing on bulk translation and TE, we conducted polysome analysis at 48 h post-reactivation; for this experiment *eIF3d* was silenced with a lentivirus vectored shRNA (as in **Figs. 2D, 2E**) and compared to a non-specific lentivirus vector control (**Fig. 3E**). The 48 h time point follows genome replication in the iSLK.219 cell model, and both late and early mRNAs are present. In our experience, there are very few polysomes detectable at 72 h post-reactivation in this model. *eIF3d* silencing resulted in a strong decrease in polysomes and an increase in monosome fractions (**Fig. 3E**). RT-qPCR analysis of monosome, light-polysome, and heavy-polysome fractions did not reveal any significant differences in TE of select viral mRNAs when eIF3d levels were reduced (**Fig. 3F**). Taken together, these data suggest that both eIF4F and eIF3d contribute to accumulation of viral proteins during lytic replication, but eIF3d depletion has a far greater effect on KSHV virion production.

**Fig. 3.**
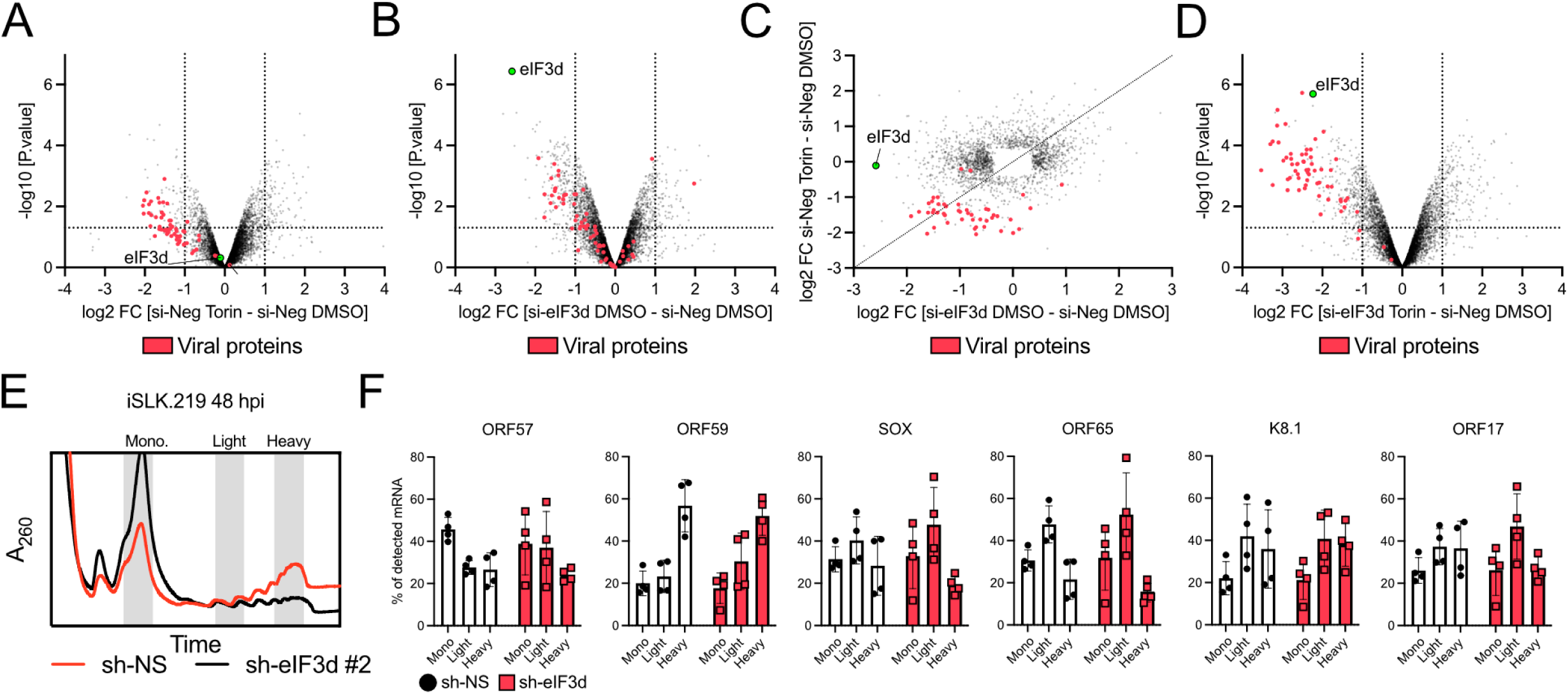
eIF4F and eIF3d both contribute to protein accumulation during lytic replication. **(A-D)** iSLK.219 cells were transfected with siRNA targeting eIF3d (si-eIF3d) or a negative control (si-Neg) and treated with doxycycline the following day. 24 h after doxycycline addition the cells were treated with 250 nM Torin of DMSO vehicle control. At 72 hpi, total protein was harvested for LC-MS/MS with data-independent acquisition (one m/z window) and label-free quantitation (n=3). Viral proteins are graphed in red and eIF3d is labelled in green. **(A)** si-Neg Torin vs si-Neg DMSO treatment demonstrating eIF4F-dependent translation. **(B)** si-eIF3d vs si-Neg demonstrating eIF3d-dependent translation. **(C)** A comparison of proteins from **(A)** and **(B)** that had a *p* value of < 0.05 in at least one of the samples. **(D)** si-eIF3d treated with Torin vs. si-Neg cells with DMSO treatment for loss of eIF3d and eIF4F. **(E-F)** iSLK.219 cells were transduced with lentiviruses encoding shRNA targeting eIF3d (sh-eIF3d) or a non-targeting (sh-NS) shRNA control. 48 h after lytic induction, cells were harvested for polysome analysis. **(E)** Polysome profile of sh-NS and sh-eIF3d. **(F)** Total RNA was extracted pooled fractions from monosomal, light polysomes, or heavy polysomes were used for RT-qPCR analysis targeting the mRNAs indicated. Data is presented as the total amount of mRNA detected in each fraction (total mRNA detected = 100%; n=4; mean ± SEM; statistical significance was determined by two-way ANOVA).

### *eIF3d* silencing reduces levels of cholesterol biosynthesis enzymes

Because eIF3d silencing caused relatively modest decreases in viral protein accumulation compared to eIF4F depletion (**Figs. 3A, 3B**), but dramatically decreased production of infectious virions by ∼4 log_10_ IU/ml (**Fig. 2B**), we hypothesized that eIF3d might specifically be required for translation of host mRNAs that are important for KSHV lytic replication. We obtained a deeper proteome of total protein harvested at 72 h post-reactivation from cells transfected with *eIF3d* siRNA compared to control siRNA using three-separate mass-charge windows. GSEA revealed that *eIF3d* silencing reduced KSHV protein levels as expected (**Figs. 4A, 4B, S7 Table**), followed by several gene sets linked to cholesterol regulation, including sterol regulatory element-binding protein (SREBP), nuclear receptor subfamily 1 group H member 3 (NR1H3) and NR1H2 (also known as Liver X Receptor α (LXRα) and LXRβ, respectively), and cholesterol biosynthesis. We annotated a custom cholesterol metabolism gene set by selecting genes from several of the down-regulated pathways which we used to annotate our volcano plots. Interestingly, the up-regulated gene sets in proteomes from *eIF3d*-deficient cells generally resembled the most down-regulated gene sets from our Torin treatment translatome, suggesting a compensatory mechanism for eIF3d insufficiency involving increased capacity for canonical eIF4F translation. This dataset stands in contrast to studies of eIF3d in human cytomegalovirus (HCMV) infection in normal human diploid fibroblasts (NHDFs), which featured regulation of unfolded protein response but no effects on cholesterol regulation ^26^. This indicates that while eIF3d can be used to support translation of host mRNA, the availability of those mRNAs can vary during virus replication, likely due to differences in host responses, host shutoff, and/or transcription. We focused on two cholesterol biosynthesis proteins that were most significantly down-regulated for further analysis: squalene epoxidase (SQLE), which is the second rate-limiting in cholesterol biosynthesis^27^, after HMG-CoA reductase (HMGCR), the target of statin drugs; and mevalonate kinase (MVK), which acts on the products generated by HMGCR **(Fig 4C)**.

**Fig. 4.**
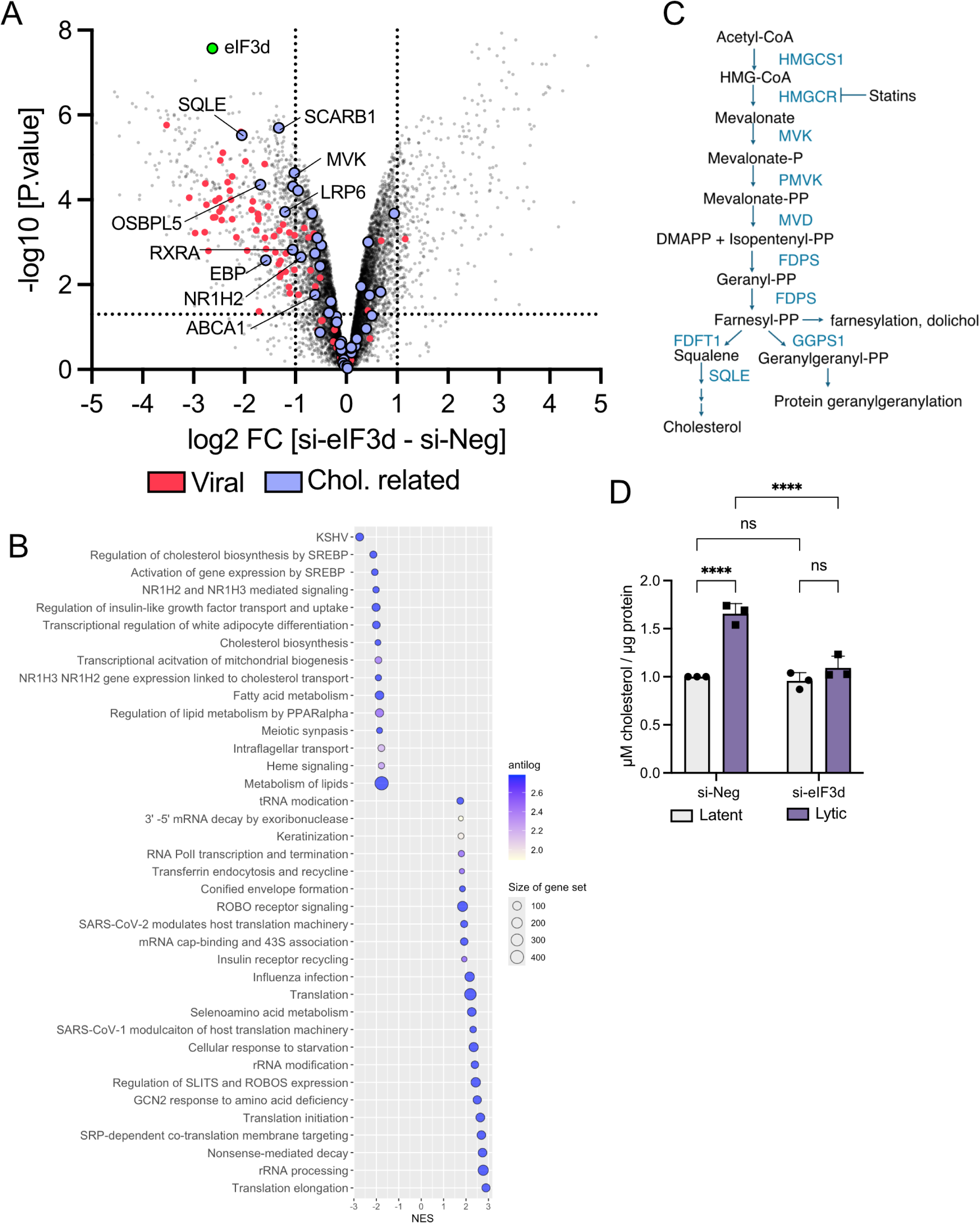
eIF3d is required for viral protein accumulation and cholesterol biosynthesis during KSHV lytic replication. iSLK.219 cells were transfected with siRNA targeting eIF3d (si-eIF3d) or a negative control (si-Neg) and treated with doxycycline the following day. **(A)** Total protein was harvested for LC-MS/MS with data-independent acquisition (three m/z windows) and label-free quantitation (n=3). **(B)** Gene set enrichment analysis with 10% false discovery rate (FDR). **(C)** General diagram of the cholesterol biosynthetic pathway. Gene names for cholesterol biosynthesis enzymes are in blue. NES = Normalized Enrichment Score. **(D)** Quantification of cholesterol abundance by luciferase-based reporter assay normalized to total protein quantity at 72 hpi (n=3; mean ± SEM; statistical significance was determined by two-way ANOVA).

To confirm the role of eIF3d in cholesterol metabolism during KSHV infection, we once again transfected iSLK.219 cells with *eIF3d* or control siRNAs prior to Dox-induced lytic reactivation. We harvested cell lysates at 72 h post-reactivation and measured total cellular cholesterol using a Cholesterol/Cholesterol Ester-Glo assay, with data normalized to total cellular protein (**Fig. 4D**). This confirmed that total cellular cholesterol significantly increased during KSHV lytic replication in an eIF3d-dependent manner. By contrast, eIF3d had no effect on the more modest levels of total cellular cholesterol found in latently infected cells. This suggests that eIF3d may be part of a response to cholesterol insufficiency, whereas cells with sufficient cholesterol maintain basal cholesterol levels without requiring eIF3d-dependent translation.

### eIF3d is required for efficient translation of SQLE and MVK and is required for accumulation of cholesterol during lytic replication

We again used proteomics to measure changes in cholesterol biosynthesis genes during lytic replication. Evaluating the changes in lytic replication compared to latent cells, there is no change in overall eIF3d abundance by 72 h post-reactivation, but there are large increases in SQLE and modest increases in HMGCS1, and MVK (**Fig. 5A, S8 Table**), suggesting that KSHV lytic replication may stimulate additional capacity in cholesterol biosynthesis. When eIF3d is depleted in latently infected cells, we see an overall decrease in the cholesterol biosynthesis proteins (**Figs. 5B, 5D, S9 Table**) but no significant change in MVK or SQLE. By contrast, *eIF3d* silencing during lytic replication markedly decreased SQLE and MVK levels (**Fig 5C**, same dataset as **Fig. 3B** with additional annotation, **S10 Table**). As an entire gene set, we see that lytic replication features a general increase in cholesterol biosynthesis enzymes, and *eIF3d* silencing depletes these proteins; more significantly, during lytic replication there is a tendency to increase cholesterol biosynthesis more so than during latency **(Fig 5D)**. Close inspection of raw intensity values confirms that *eIF3d* silencing prevents the upregulation of SQLE and MVK that accompanies lytic replication **(Fig. 5E)**. Using RNA isolated from previous polysome profiling experiments **(Figs. 3E-F)**, we used RT-qPCR to demonstrate that *eIF3d* silencing reduces translational efficiency of *SQLE* and *MVK*, moving these mRNAs from heavy into light polysome fractions, whereas mRNAs encoding HMGCS and low-density lipoprotein receptor (LDLR) were relatively unaffected **(Fig. 5F)**, consistent with their unchanged total protein abundance in eIF3d deficient cells. These data suggest that eIF3d is required for translation of mRNAs encoding SQLE and MVK during lytic replication, and downstream increases in total intracellular cholesterol.

**Fig 5.**
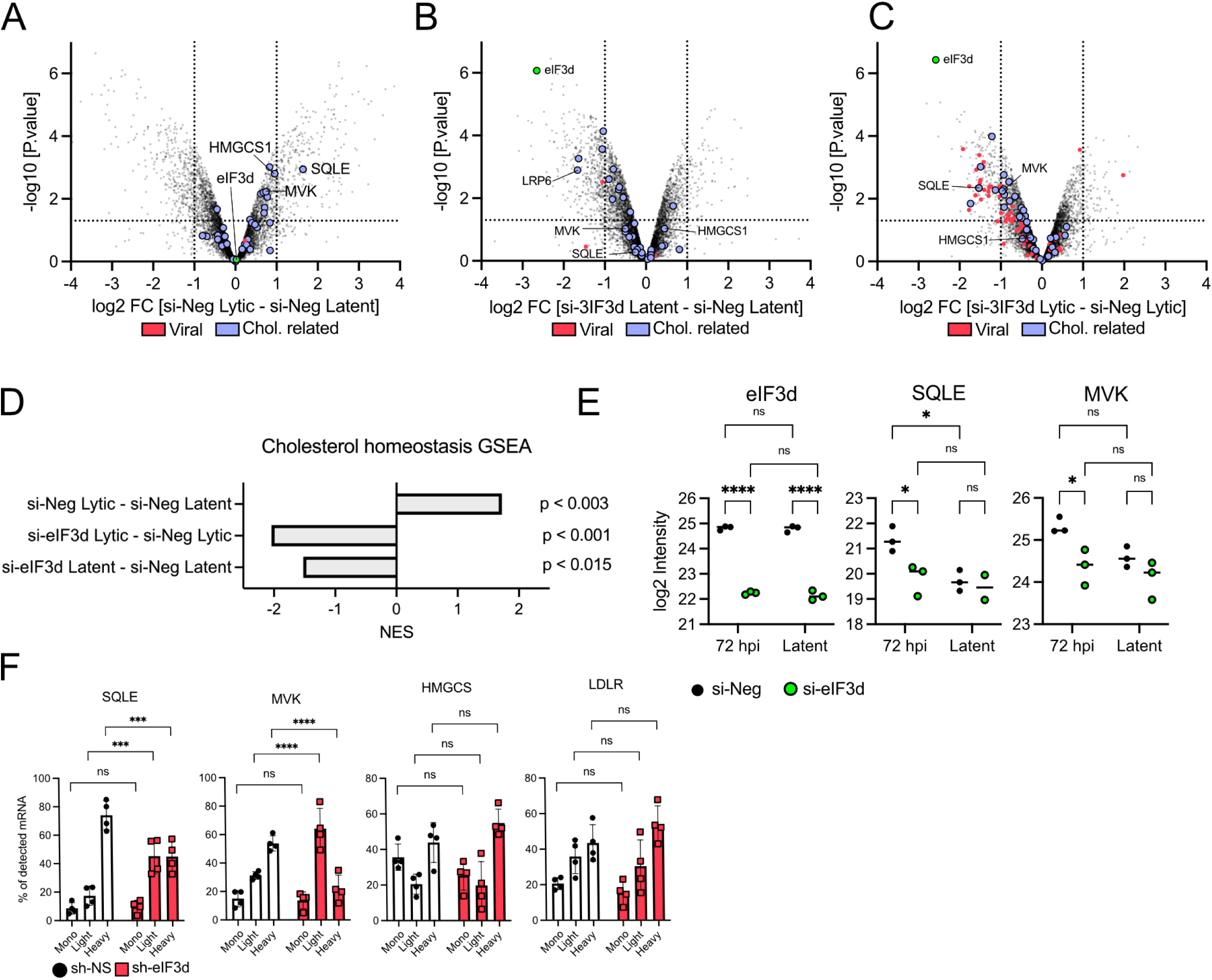
eIF3d is required for efficient translation and accumulation of SQLE and MVK during KSHV lytic replication. **(A-C)** iSLK.219 cells were transfected with siRNA targeting eIF3d (si-eIF3d) or a negative control (si-Neg) and treated with doxycycline the following day. 24 h after doxycycline addition the cells were treated with 250 nM Torin of DMSO vehicle control. At 72 hpi, total protein was harvested for LC-MS/MS with data-independent acquisition (one m/z window) and label-free quantitation (n=3). Viral proteins are graphed in red; proteins related to cholesterol biosynthesis are in blue; eIF3d, SQLE, and MVK are labelled individually in green. **(A)** si-Neg Lytic vs. si-Neg Latent for changes associated with lytic replication. **(B)** si-eIF3d vs. si-Neg during latency for eIF3d-dependence in latently infected cells. **(C)** si-eIF3d vs. si-Neg in lytic cells in matched samples of **(A)** and **(B)**. **(D)** Cholesterol biosynthesis Gene set enrichment analysis (GSEA) for **(A-C)** NES = Normalized Enrichment Score. **(E)** log2 transformed intensity value for eIF3d, SQLE, and MVK as measured for values represented in **(A-C)**. **(F)** Polysome analysis of cholesterol biosynthesis genes as in Fig. 3E-F. (total mRNA detected = 100%; n=3; mean ± SEM; statistical significance was determined by two-way ANOVA).

### eIF3d is required for increased SQLE and LDLR accumulation in response to cholesterol scarcity

It was recently demonstrated that in cancer cells, when eIF4F is unavailable, eIF3d reprograms cholesterol homeostasis, inducing alternative lipid uptake and promoting cholesterol transport ^28^. This response required LRP6, which we also observed to be decreased upon eIF3d depletion **(Fig. 4A)**. While this supports a role for eIF3d in cholesterol uptake and transport, our data also suggested an additional role in cholesterol biosynthesis. We reasoned that eIF3d-dependent control of cholesterol biosynthesis might be more apparent under conditions that increase demand, such as growth in cholesterol-depleted serum. To test this directly, we silenced *eIF3d* in uninfected iSLK epithelial cells and evaluated the contribution of eIF3d to the cellular response to cholesterol scarcity by label-free proteomics. Incubating control siRNA transfected iSLK cells in media formulated with lipoprotein-depleted serum (LPDS) for 48 h caused a significant increase in steady-state levels of enzymes of the mevalonate pathway (e.g. HMGCR, farnesyl-diphosphate farnesyltransferase 1 (FDFT1, also known as squalene synthase), SQLE) as well as the LDLR protein responsible for cholesterol uptake; by contrast, the cholesterol efflux transporter ATP Binding Cassette Subfamily A Member 1 (ABCA1) ^29^ was downregulated (**Fig. 6A, S11 Table**). This indicates that iSLK cells respond to cholesterol scarcity by attempting to increase cholesterol biosynthesis and uptake, while reducing efflux, as expected. However, when *eIF3d* was silenced, FDFT1, SQLE and LDLR upregulation was muted during cholesterol scarcity; HMGCR upregulation was eIF3d-independent (**Fig. 6B, S12 Table**). *eIF3d* silencing in uninfected cells cultured in rich medium caused strong down-regulation of LRP6 and reduced SQLE abundance, with a much larger effect on SQLE than observed in latently infected cells; ABCA1 was also strongly decreased **(Fig 6C, S13 Table)**. Together, these changes suggest that *eIF3d* silencing disrupts multiple arms of cholesterol homeostasis, including uptake, biosynthesis, and efflux. The decrease in ABCA1 is consistent with the possibility that cells limit cholesterol export when cholesterol biosynthesis and import are impaired. Examination of intensity values demonstrate the eIF3d dependence of SQLE and LDLR accumulation, and eIF3d-independent regulation of HMGCR (**Fig. 6D**). Taken together with the requirement for eIF3d to support cholesterol accumulation in lytic cells, these data suggest that eIF3d supports cholesterol homeostasis by promoting SQLE accumulation in response to cholesterol scarcity.

**Fig 6.**
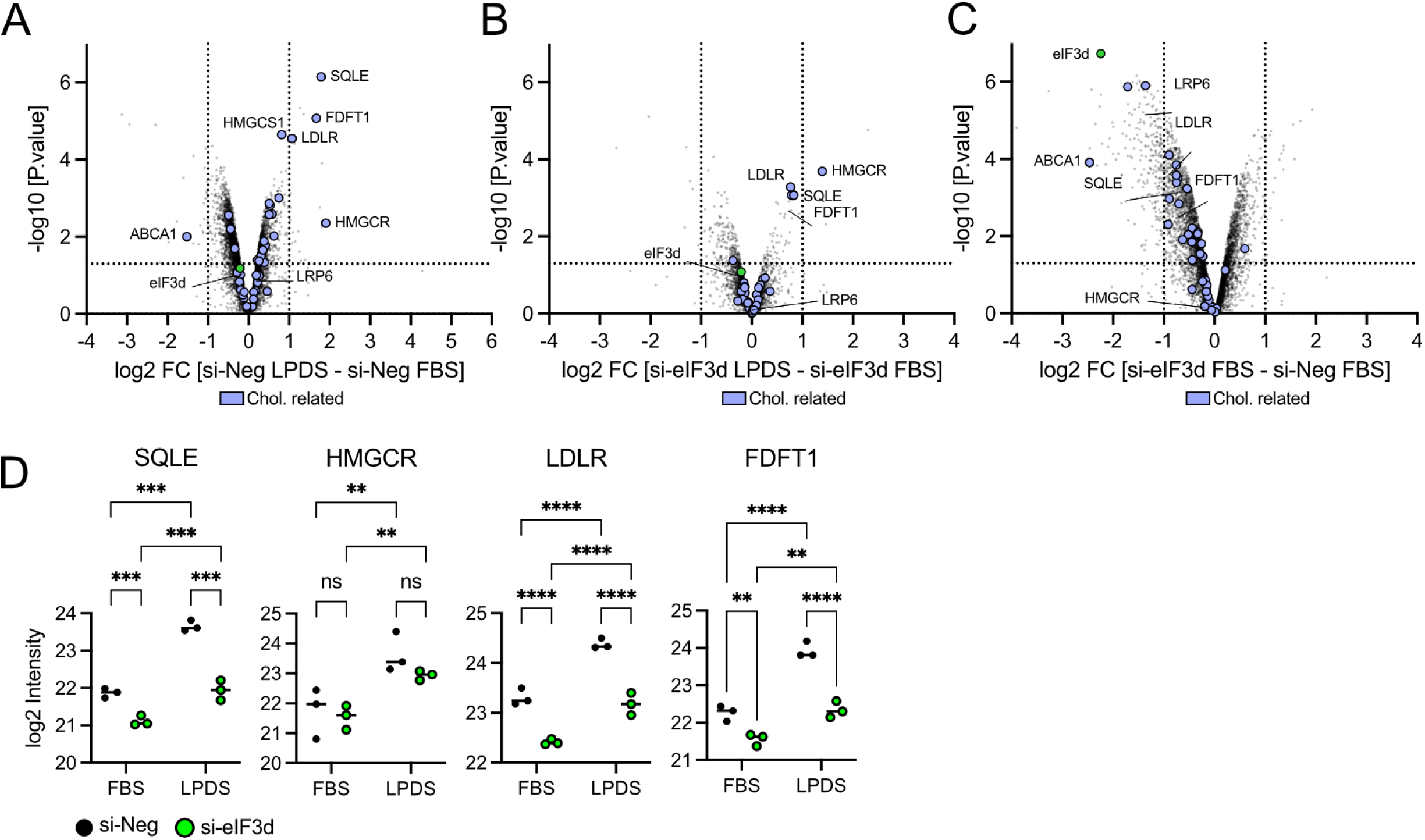
eIF3d is required for increased SQLE expression during cholesterol starvation. **(A-C)** Uninfected iSLK cells were transfected with siRNA targeting eIF3d (si-eIF3d) or a negative control (si-Neg). The following day the cells were washed with PBS twice and medium was replaced with DMEM containing fetal bovine serum (FBS) lipoprotein-depleted serum (LPDS). After 48h of cholesterol starvation, total protein was harvested for LC-MS/MS with data-independent acquisition (one m/z window) and label-free quantitation (n=3). Proteins related to cholesterol biosynthesis are in blue; eIF3d, SQLE, HMCGR, and LDLR are labelled individually in green. **(A)** si-Neg LPDS vs. si-Neg FBS for changes associated with cholesterol starvation. **(B)** si-eIF3d LPDS vs. si-eIF3d FBS during latency for eIF3d-dependence in latently infected cells. **(C)** si-eIF3d vs. si-Neg in lytic cells in matched samples of **(A)** and **(B)**. **(D)** log2 transformed intensity value for eIF3d, SQLE, and MVK as measured for values represented in **(A-C)**. **(E)** Polysome analysis of cholesterol biosynthesis genes as in Fig. 3E**-F**. (total mRNA detected = 100%; n=4; mean ± SEM; statistical significance was determined by two-way ANOVA). **(G)** Quantification of cholesterol abundance by luciferase-based reporter assay normalized to total protein quantity at 72 hpi (n=3; mean ± SEM; statistical significance was determined by two-way ANOVA).

### Productive KSHV lytic replication requires access to cholesterol through biosynthesis or uptake

We demonstrated that eIF3d is required to support KSHV lytic replication and promotes the accumulation of key proteins involved in cholesterol biosynthesis (e.g. MVK, FDFT1, SQLE) and uptake (LRP6), suggesting that lytic replication depends on intracellular cholesterol. Our results suggest that when eIF3d is depleted, cholesterol cannot accumulate normally during lytic replication. We find that eIF3d depletion reduces the expression of key cholesterol biosynthetic proteins, and previously published work demonstrates that eIF3d inhibition limits uptake of cholesterol from lipoproteins ^28^. However, it remained unclear to what extent cholesterol is required to complete lytic replication, and whether lytically infected cells are competent to synthesize or import cholesterol. To test this, we used the selective SQLE inhibitor NB-598 to inhibit cholesterol biosynthesis, and LPDS to limit lipoprotein-derived cholesterol uptake. The inhibitor was well tolerated for 48 h of treatment in latent or lytic cells grown in medium with full FBS or LPDS **(Figs. 7A, 7B)**. LPDS starvation during lytic replication was associated with reduced viability compared to lytic cells in full medium **(Fig. 7C)**. We observed no difference in release of infectious virions from cells in full FBS medium or LPDS, suggesting that lipoprotein-derived cholesterol uptake is not strictly required for virion production **(Fig. 7D)**. Treatment of cells with NB-598 at concentrations up to 5 µM also had little effect, suggesting that cholesterol biosynthesis is similarly not strictly required for virion production **(Fig. 7E)**. Importantly, when we combined SQLE inhibition with LPDS starvation, there was a strong defect in virion production that extended to the lowest concentration of inhibitor tested. By contrast, the farnesyltransferase inhibitor lonafarnib had a limited effect on virion release, indicating that protein farnesylation, a separate downstream product of the mevalonate pathway, is dispensable for productive KSHV replication **(Figs. 7F-H)**. These results suggest that additional intracellular cholesterol required to support KSHV lytic replication can be supplied by either biosynthesis or import, but that at least one of these mechanisms must be available. Taken together with the requirement for eIF3d in cholesterol import and cholesterol biosynthesis, these findings suggest that the specific requirement for eIF3d in KSHV virion production is at least in part related to cholesterol metabolism.

**Fig. 7.**
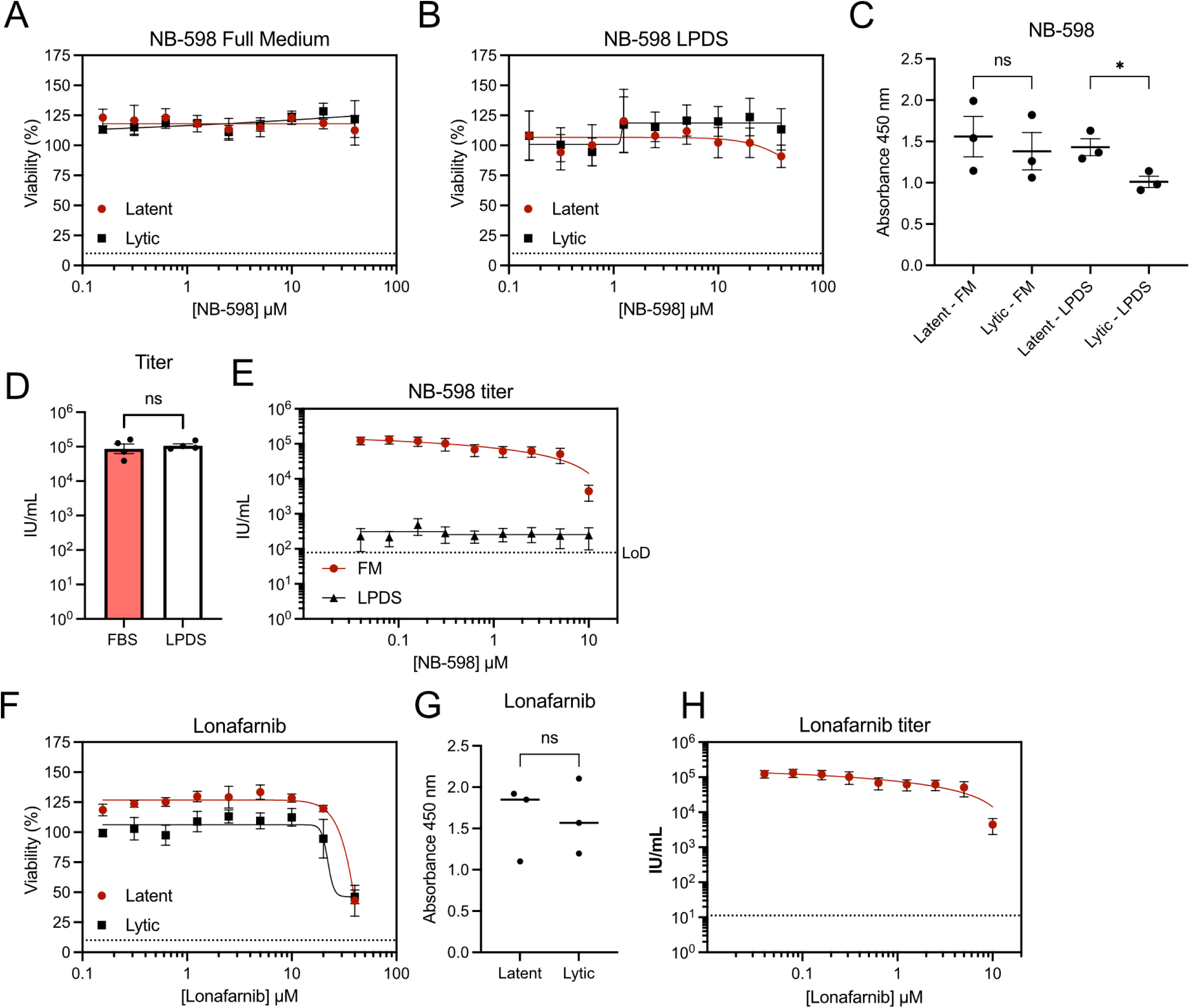
Restricting cholesterol biosynthesis and uptake inhibits KSHV virion production. **(A)** Latently infected or lytically replicating iSLK.219 cells treated with varying concentrations of SQLE inhibitor NB-598. Viability was determined at 48h using colorimetric CCK-8 assay. **(B)** As in **(A)** but with lipoprotein-depleted serum (LPDS). **(C)** raw intensity values of DMSO treated samples from **(A)** and **(B)** (n=3; mean ± SEM; statistical significance was determined by one-way ANOVA) titer at 96hpi from iSLK.219 cells treated with in FBS-containing medium or LPDS (n=4; mean ± SEM; statistical significance was determined by paired T-test). **(E)** As in **(D)** with varying concentrations of NB-598 (n=4; mean ± SEM). **(F)** Cell viability was determined as in (A) using the FTase inhibitor Lonafarnib **(G)** raw intensity values of DMSO treated samples from **(F)** (n=3; mean ± SEM; statistical significance was determined by paired *t*-test). **(H)** as in **(E)** in FBS-containing medium.

### eIF3d silencing is associated with REDD1 downregulation and increased mTORC1 signaling

When eIF4F assembly is inhibited in response to nutrient scarcity or treatment with mTORC1 inhibitors, residual translation can be supported by eIF3d ^5^. Consistent with this, inhibition of mTORC1 appears to be sufficient to promote eIF3d-dependent translation ^12^. Cholesterol is also important to maintain mTORC1 association with the lysosome and support its activation ^30,31^. Since eIF3d appeared to influence cholesterol metabolism, we asked whether eIF3d silencing might impair mTORC1 activation. In our data, GSEA of the translatome during mTORC1 inhibition showed down-regulation of several pathways related to translation, including initiation, elongation, and nonsense-mediated decay (**Fig. 1E**). These were the same pathways that were up-regulated in the eIF3d-silencing proteomes (**Fig. 4B**), leading us to ask whether eIF3d silencing might influence mTORC1 activation. By reanalyzing our data from *eIF3d* silencing of lytic iSLK.219 cells at 48 h post-reactivation, we identified a small group of proteins involved in mTOR regulation that accumulate to higher levels (e.g. WD Repeat Domain 24 (WDR24), and Ras-Related GTP-Binding Protein B (RRAGB, also known as RagB)) or lower levels (e.g. Protein Regulated In Development And DNA Damage Response 1 (REDD1, also known as DNA Damage-Inducible Transcript 4 Protein (DDIT4)), and cytosolic arginine sensor for mTORC1 subunit 2 (CASTOR2)) in the absence of *eIF3d* (**Fig. 8A**) ^32^. Amongst these proteins, we noted the ∼30-fold downregulation of REDD1, an important negative regulator of mTORC1 signaling during hypoxia ^33^. Normally, REDD1 inhibits mTORC1 indirectly by promoting TSC1/TSC2 complex activity; TSC2 is a GTPase-activating protein that stimulates RHEB-GTP hydrolysis, lowering the amount of active RHEB-GTP available to allosterically activate mTORC1 at the lysosome ^31^. To test the activation state of mTORC1, we silenced *eIF3d* in medium containing LPDS and provided exogenous cholesterol in the form of LDL. We determined that *eIF3d* silencing prevented REDD1 accumulation in latently KSHV infected iSLK.219 cells, lytically infected iSLK.219 cells, and in uninfected iSLK cells (**Fig. 8B**). In the latent and lytic cells, reduced REDD1 correlated with increased phosphorylation of ribosomal protein S6 (S6), consistent with increased mTORC1-S6K signaling. We did not observe a change in mTORC1 activation in uninfected cells, possibly because LPDS produces less severe cholesterol limitation than acute cholesterol extraction with methyl-β-cyclodextrin ^30^. This may indicate that mTORC1 signaling in infected cells is more sensitive to cholesterol limitation. Together, these data suggest that when eIF3d is unavailable, cells may de-repress mTORC1 signaling to support efficient eIF4F assembly.

**Fig. 8.**
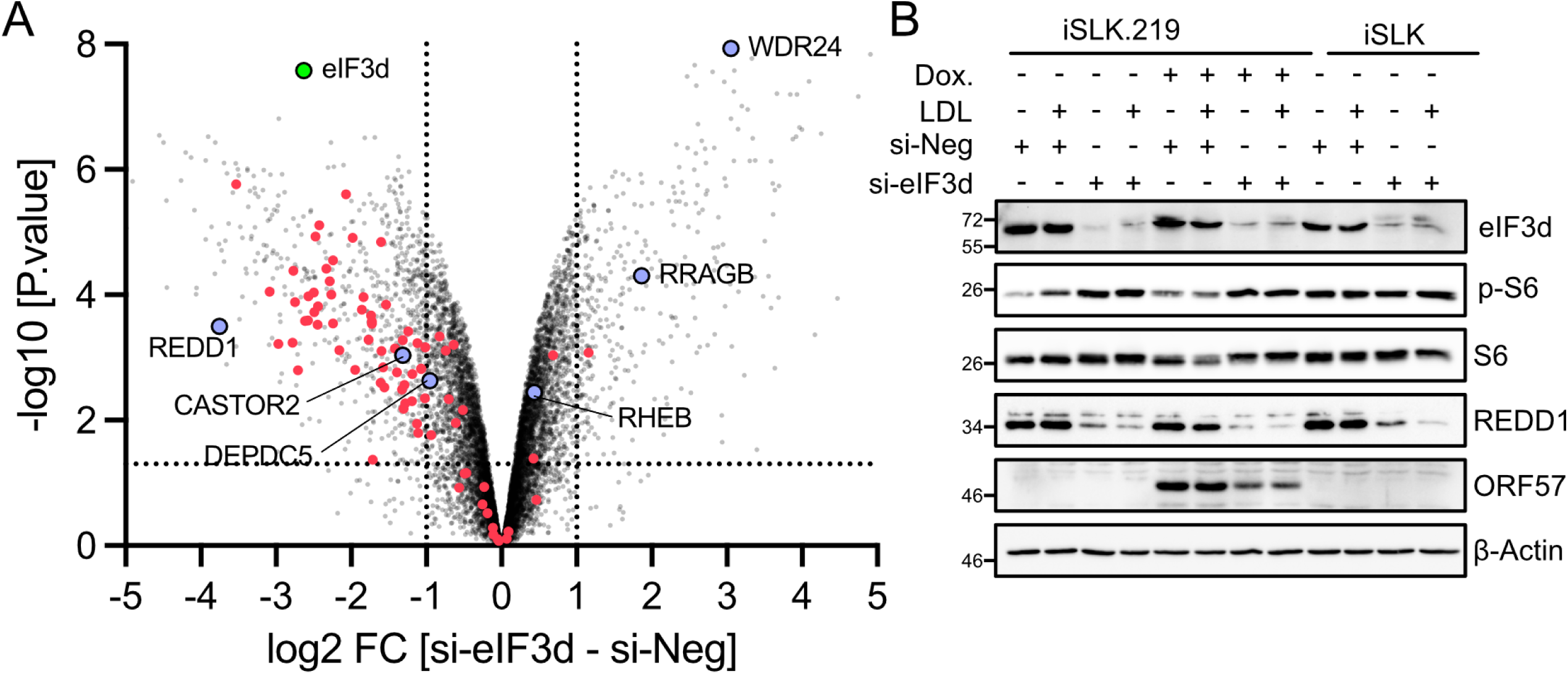
mTORC1 remains active during cholesterol starvation in eIF3d KD. **(A)** Recreated from **Fig 4A** with mTORC1 regulatory proteins labelled in blue. **(B)** Western blot of whole cell lysates of uninfected, latently infected, or 48 h post-doxycycline (Dox.) treatment (hpi) previously transfected with siRNA targeting eIF3d (si-eIF3d) or a negative control siRNA (si-Neg) in LPDS-containing medium. 100 µg/mL of LDL was added to samples as indicated for 1 h prior to harvest.

## DISCUSSION

KSHV mRNAs broadly resemble their cellular counterparts, which should allow their recruitment to ribosomes in an eIF4F-dependent manner. Torin treatment during lytic replication depletes eIF4F and shifts TOP-containing mRNAs encoding ribosomal proteins and TIFs into monosomal fractions ^6^; eIF3d knockdown also depletes heavy polysomes during lytic replication. Heavily translated polysome fractions contain viral mRNA, which remains when either eIF4F or eIF3d is depleted. Despite little difference in the efficiency of translation of viral mRNAs in response to Torin treatment **(Fig. 1D)** or *eIF3d* silencing **(Fig. 3F)**, there is decreased accumulation of viral proteins (**Fig. 3**). Thus, we speculate that in this well-established KSHV infection model, viral mRNA is not specifically regulated or dependent on either of these initiation systems and can possibly use either with similar efficiency, but the availably of these systems is important to optimize viral protein synthesis throughout the relatively slow-moving KSHV lytic cycle. The additive effects of Torin treatment and eIF3d knockdown on viral protein accumulation are consistent with this model (**Fig. 3C**).

KSHV appears to access multiple translation initiation systems to maximize protein production. KSHV encodes several proteins that enforce mTORC1 activation, which supports eIF4F formation ^34^. Our data suggest that the virus also activates eIF3d-dependent translation, but the underlying mechanism is not currently known. Glucose starvation has been shown to allow access to the cap-binding pocket of eIF3d ^13^, and KSHV might use a similar metabolic signaling pathway or exploit another yet to be described regulatory pathway to access eIF3d. HIF2⍰ is stabilized in normoxia during lytic replication by ORF34 ^35^. This likely permits HIF2⍰ to stimulate formation of the eIF4F_H_ hypoxic translation initiation complex in normoxia during KSHV lytic replication, which would make it available to support translation of viral mRNA ^36^. eIF4F_H_ and eIF4F are also known to co-exist in cells under physiological oxygen tensions ^37^. Because we see decreased accumulation of viral mRNA with eIF4F disassembly and eIF3d silencing, it may be that eIF4F_H_ is not solely responsible for the translation of viral mRNA and may instead act in parallel with eIF4F and eIF3d. The accumulating evidence suggests that KSHV evolved to opportunistically use any available translation initiation machinery to translate viral mRNA.

In contrast to the potentially generalist nature of KSHV mRNA, some cellular mRNAs are tightly regulated by specific initiation systems. mTORC1 and LARP1 tightly regulate TOP containing mRNA ^6,11,38^. eIF3d has been shown to have specific client mRNAs, such as the well characterized *c-Jun* and *Raptor* mRNAs that contain a long and structured 5□UTR, which excludes initiation by eIF4F but is compatible with eIF3d cap-binding ^13,14^. The 5□-UTRs of KSHV mRNA are generally short compared to these cellular 5□-UTRs and we think that feature might help their access to different translation systems. We found that eIF3d is required for efficient translation of SQLE and MVK mRNA during lytic replication (**Fig. 5F**). Neither SQLE or MVK mRNA was identified as being an “eIF3d-specialized” mRNA when eIF4E was inhibited ^14^. SQLE was detected as binding eIF3d in a pulldown assay ^13^ during glucose starvation and chronic ER stress ^39^, which further supports a role for eIF3d in SQLE translation. SQLE and MVK are not regulated translationally by eIF3d during HCMV infection ^26^, similar to what we observed in cells latently infected with KSHV (**Fig. 5B**). This suggests that while eIF3d might be required for enhanced cholesterol biosynthesis, it is not sufficient and likely requires support from an appropriate transcriptional program.

It is currently unclear how cholesterol regulates KSHV lytic replication, or how cholesterol is regulated during KSHV latency, although from our data it is clear cholesterol must be synthesized or imported. Interestingly, HIV infected individuals who are also taking statins have a 51% reduced risk of developing Kaposi’s sarcoma ^40^, suggesting that cholesterol dependency might be a therapeutically targetable vulnerability of KSHV. Herpesvirus egress requires the nascent capsids in the nucleus to pass through several membranes, including the inner and outer nuclear membranes during primary envelopment and *trans*-Golgi network membranes during secondary envelopment^41^. KSHV secondary envelopment has been shown to be inhibited by extracting cholesterol from the cell with methyl-β-cyclodextrin ^41^. In general, the amount of cholesterol in membranes other than the plasma membrane is relatively low, but these lower amounts of cholesterol at internal membranes likely have important roles in membrane fluidity and lipid organisation. We think it is likely that reduced cholesterol availability when eIF3d is depleted affects the ability of KSHV capsids to access or manipulate these membranes for egress. During latency, KSHV alters host cell metabolism by stimulating the Warburg effect and *de novo* fatty acid synthesis ^42,43^. KSHV microRNAs expressed during latency limit expression of key genes in the cholesterol biosynthesis pathway ^44^. Inhibition of these metabolic adaptations with glycolysis or fatty acid synthesis inhibitors leads to cell death ^42,43^. Because eIF3d supported the induction of select cholesterol biosynthetic enzymes during cholesterol scarcity in uninfected cells, these findings suggest that KSHV may access a normal cellular adaptive pathway to help meet the cholesterol demands of lytic replication. We speculate that cholesterol is likely depleted or limited in latently infected cells and this defect must be rapidly corrected to support the cholesterol requirement of lytic replication. Taken together, our data suggest a model where KSHV mRNA is efficiently translated by several different translation initiation complexes and the virus endeavours to activate several of these to maximize protein production. Reactivation from latency though comes with some specific challenges where cholesterol metabolism must be rapidly engaged and this requires eIF3d-dependent translation. In this way, KSHV is dependent on eIF3d not to translate its own mRNA, but to instead support the translation of essential host mRNAs.

## METHODS

### Cells

Human embryonic kidney (HEK) 293T and 293A cells, as well as iSLK and iSLK.219 cells (a kind gift from Don Ganem ^22^), were cultured in Dulbecco’s modified Eagle’s medium (DMEM; ThermoFisher, 11965118) supplemented with heat-inactivated 10% fetal bovine serum (FBS, ThermoFisher, A31607-01), 2 mM L-Glutamine (ThermoFisher, 25030081), and 100 U/mL penicillin and 100 µg/mL streptomycin (ThermoFisher, 15140122). For experiments with lipoprotein-depleted serum (LPDS; Kalen Biomedical #800100-1), serum was added for DMEM at 10% v/v. All cells were maintained at 37^°^C in 5% CO_2_ atmosphere. iSLK.219 cells were cultured with 10 mM puromycin (ThermoFisher, A1113803) to maintain episome copy number of rKSHV.219; puromycin was omitted from cells seeded for experiments. To induce lytic replication via expression of the RTA transgene in iSLK.219 cells, 1 μg/mL of doxycycline (dox; Sigma, D9891) was added to the cells.

### Chemicals

Torin-1 ^21^, referred to as “Torin” here (Toronto Research Chemicals) was resuspended in DMSO, which was used as a vehicle control at 0.1% (vol/vol) in all experiments. Unless otherwise stated, Torin was used at a concentration of 250nM. SQLE inhibitor NB-598 (Cayman Chemicals #14912) and Lornafarnib (Sigma #SML1457) were diluted in DMSO. Human LDL was added to cells at a concentration of 100 µg/mL (Lee Biosolutions # 360-10).

### siRNA

iSLK.219 cells were detached with 0.05% Trypsin-EDTA (ThermoFisher, 2530054), resuspended in antibiotic-free medium, and transfected with siRNA targeting *eIF3d* (ThermoFisher AM16708, Assay ID: 13732) or the negative control siRNA (ThermoFisher, Neg. Control #2, 4390846) using Lipofectamine RNAiMAX (ThermoFisher, 13778075) in Opti-MEM I (ThermoFisher, 31985070) as per the manufacturer’s instructions. Transfection complexes were mixed with cells in suspension and added to plates.

### shRNA gene silencing by transduction in suspension

Lentiviruses derived from pLKO were derived from transfection of 293T cells with packaging plasmids pMD2.G and psPAX2 (gifts from Didier Trono; Addgene plasmids # 12259 and # 12260) using 1 mg/mL polyethylenimine MAX (Polysciences #24765) in water. To transduce cells, iSLK.219 cells were detached with trypsin, counted, and mixed with 1:20 dilution of lentivirus and 4 µg/mL polybrene (hexadimethrine bromide, Sigma H9268) in suspension in a small volume (∼2mL for each 15cm plate seeded) for 15 minutes, with frequent inversion to mix. Cells were then added to a 15 cm dish and medium was topped up to 20 mL. shRNA sequences for *eIF3d* were selected from the RNAi Consortium (TRCN0000156633); pLKO vectors were generated according to the RNAi Consortium shRNA cloning protocol (https://portals.broadinstitute.org/gpp/public/resources/protocols). A non-targeting shRNA sequence was used as a control in all experiments, pLKO-NS-BSD (a gift from Keith Mostov; Addgene plasmid # 26655 ^45^). Transduced cells were freshly generated for every experiment; we could not derive iSLK.219 cells with stably knocked down *eIF3d* and found that the cells would not re-adhere on passage. Similarly, we could not select cells and re-seed them at densities appropriate for polysome analysis. Thus, the transduced cells were not selected with blasticidin, and we instead relied on a low dilution/high MOI of lentivector to effectively transduce the population and obtain a good knockdown.

### Cell viability

Cells were seeded into 96-well tissue culture plates in 100 µL medium per well. The next day, media was replaced with fresh media and cerivastatin at the indicated concentrations; to measure the effects of cerivastatin on cell viability during lytic replication, this media change included 1 µg/mL doxycycline. Cells were left to incubate with cerivastatin for 48 h prior to addition of 10 µL of Cell Counting Kit 8 (CCK-8; ApexBio, K1018). Only the inner wells were used to reduce variation from evaporation, and samples were seeded in triplicate. We chose to use a colourimetric assay instead of a fluorescence-based viability assay to avoid any potential interference from the bright GFP and RFP expression from the iSLK.219 cells. After 1 h of incubation with CCK-8, absorbance was read at 450 nm with correction at 600 nm using a CLARIOstar Plus instrument (BMG Labtech). Values were imported into Prism (v10.6; GraphPad) and 50% cytotoxic concentration (CC50) values were calculated by fitting a non-linear curve to the data. The values from the 10, 5, and 2.5 µM cerivastatin treatments were separately analysed by two-way ANOVA using Prism.

### Western blotting

iSLK.219 cells were washed once with ice-cold PBS then lysed with 2x Laemmli buffer (4% [wt/vol] sodium dodecyl sulfate (SDS), 20% [vol/vol] glycerol, 120 mM Tris-HCl [pH 6.8]) directly in the well. DNA was sheared by repeated pipetting with a 21-gauge needle, or by centrifugation through a QIAshredder column (Qiagen, 79656) at 13,000xg for 1 min, prior to 100 mM dithiothreitol (DTT) addition and boiling at 95^°^C for 5 min. Samples were stored at -20^°^C until analysis. Total protein concentration was determined by DC Protein Assay (Bio-Rad, 5000116) against a bovine serum albumin (BSA) (Bioshop, ALB001.25) standard curve and measured in a 96-well plate format at 750 nm using an Eon (BioTek) microplate spectrophotometer. Equal quantities of 10 μg total protein were loaded in each well of a polyacrylamide gel and subjected to SDS-PAGE. Proteins were transferred to polyvinylidene difluoride (PVDF) membranes using the Trans-Blot Turbo RTA Midi 0.2 μM PVDF Transfer Kit (Bio-Rad, 1704273) and a Trans-Blot Turbo Transfer System (Bio-Rad). Membranes were blocked with 5% BSA in Tris-buffered saline, 0.1% [vol/vol] Tween-20 (TBS-T) before probing overnight at 4^°^C with the following antibodies at 1:1000 dilution: 4E-BP1 (New England Biolabs Canada (NEB), #9644), β-actin (NEB #5125 or NEB #4970), Rps6 (S6; NEB #2217) phosphoSer235/6-Rps6 (NEB #4858), ORF57 (Santa Cruz Biotechnologies sc-135746), LANA (a kind gift of Don Ganem) and REDD1 (ProteinTech #10638-1-AP). Membranes were washed with TBS-T and incubated with HRP-linked secondary antibodies diluted 1:3000 in 5% BSA in TBS-T for 1 h at room temperature. Secondary antibodies used: anti-rabbit, HRP-linked (NEB # 7074, 1:3000 dilution), and anti-mouse, HRP-linked (NEB, #7076). Immunoblots were developed with Clarity ECL reagent (Bio-Rad, 170-5061) or Clarity Max ECL reagent (Bio-Rad, 170-5062) and imaged on a ChemiDoc MP Imaging System (Bio-Rad). Molecular weights were determined using the Broad Range, Color Prestained Protein Standards (NEB, P7719S). Molecular weights in kDa are indicated on the left of blot images. Images were analyzed using Image Lab 6.1 (Bio-Rad). Images were cropped and annotated using Affinity Designer (Serif). See **S1_raw_images** for uncropped and unadjusted images underlying all western blot data.

### Virus titering

iSLK.219 cells were seeded in a 96-well plate. In some experiments, these cells were transfected with siRNAs as described above during the plating procedure. The following day, the supernatant was replaced with fresh medium containing 1 µg/mL doxycycline to induce reactivation from latency. For experiments where Torin was added, the drug was spiked in 24 h after doxycycline addition (**Fig. 2A**); serial dilutions of cerivastatin were added at the time of doxycycline addition (**Fig. 6C**). Cell supernatants were harvested at 96 h post-reactivation, diluted 1/20 and then used to infect 293A cells by spinoculation, whereby after the inoculum was added, plates were centrifuged at 800 x *g* at 37^°^C for 90 min, then moved to an incubator before fixation 24 h later. Infected cells were counterstained with Hoechst 33342 (ThermoFisher, 62249) to label DNA-containing nuclei and GFP+ cells were counted using an automated fluorescent microscope and Cell Profiler ^46^ or Imaris Microscopy analysis software [Oxford Instruments]. The IU/mL was determined by counting the number of uninfected cells and calculating the corresponding MOI.

### RT-qPCR

Total RNA from cells was extracted using the RNeasy Plus Mini Kit (Qiagen, 74134) following the manufacturer’s protocol. Synthesis of cDNA was performed using the Maxima H Minus First Strand cDNA Synthesis Kit (ThermoFisher Scientific, K1652) using random hexamer primers. Quantitative PCRs were performed using 200 nM (final) primers, 1:250 (final) diluted cDNA, and 1X GoTaq qPCR Master Mix (Promega, A6002). Amplifications were performed using a CFX Connect Real-Time PCR Detection System (Bio-Rad) and Bio-Rad CFX Manager 3.1 software. Primer sequences are listed in **Table 1**. Due to common usage of polyA signals, qRT-PCR primers could amplify mRNA originating from more than one viral promoter ^2^. For simplicity, PCR products are referred to by the mRNA coding region targeted by the primers.

**Table 1.**
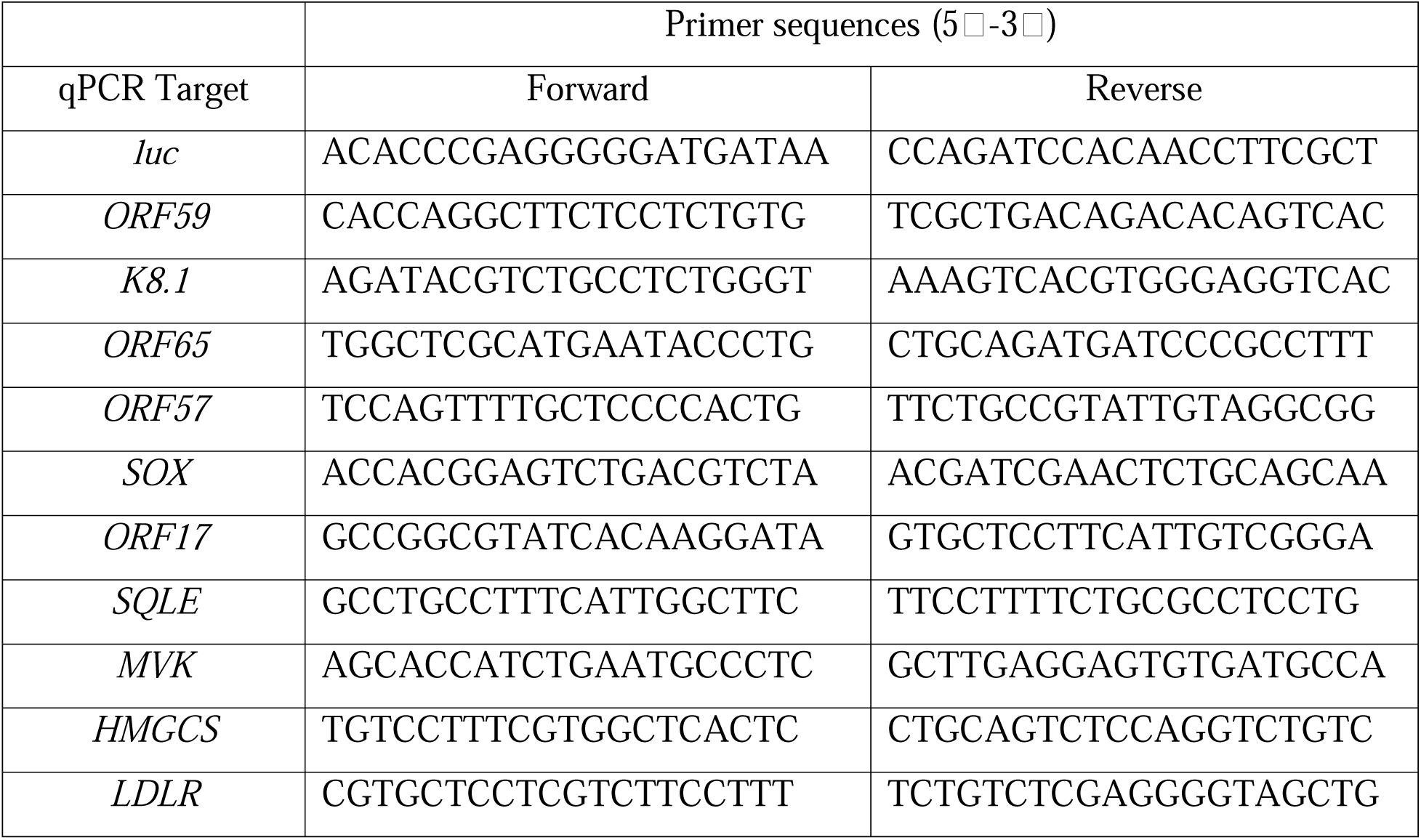
Oligonucleotide sequences.

### Cholesterol quantification

Cholesterol quantity was measured using the Cholesterol/Cholesterol Ester-Glo assay (Promega, J3190) essentially as per the manufacturers’ directions. iSLK.219 cells transfected with siRNA and seeded in a 96 well plate in triplicate. The following day the medium was refreshed and 1 µg/mL doxycycline was added to stimulate lytic replication. After 72 h the supernatant was removed, the cells were washed twice with ice-cold PBS and then lysed in 50 µL of the Cholesterol lysis solution and the cells were incubated for 30 min at 37^°^C. 10 µL of the lysis solution was transferred to a white 96-well plate and mixed with 10 µL of cholesterol detection reagent with esterase. The plate was then incubated at room temperature for 1 h and luminescence was measured on a CLARIOstar Plus plate reader. To normalize the cholesterol measurements to the number of cells in the plate, a DC protein assay was performed as described above.

### Polysome Isolation

Polysomes were isolated by ultracentrifugation of cytosolic lysate through a 7-47% wt/vol linear sucrose gradient (20 mM Tris HCl, 300 mM NaCl, 25 mM MgCl_2_ in DEPC-treated or nuclease-free water) lysis buffer with RNase and protease inhibitors, as described in ^47^. High salt conditions were used to isolate RNA from gradients and low salt conditions were used for isolating proteins. Gradients were prepared using manufactures’ settings on a Gradient Master 108 (Biocomp). For each gradient, ∼8 x 10^6^ iSLK.219 cells were used. Cells treated with 100 µg/mL cycloheximide (CHX, Sigma C1988) for 3 min prior to harvest. Cells were washed with ice-cold phosphate buffered saline (PBS) and scraped in a tube containing the spent media and PBS wash. The cells were pelleted by centrifugation for 5 min at 500 x *g* and washed again with ice-cold PBS. Cell pellets were resuspended in lysis buffer (20 mM Tris HCl, 300 mM NaCl, 25 mM MgCl_2_, with 1% v/v Triton X-100, 400 units/ml RNAseOUT (Invitrogen), 100 µg/mL CHX, and cOmplete protease inhibitors (Roche, 04693116001) for 10 min on ice. Lysates were centrifuged for 10 min at 15,000 x *g*. The supernatant was overlaid on sucrose gradients. Gradients were centrifuged at 39,000 rpm for 90 min on a SW-41 rotor. The bottom of the centrifuge tube was punctured and 60% wt/vol sucrose was underlaid by syringe pump to collect 500 µL fractions from the top of the gradient with simultaneous A_260_ measurement using a UA-6 detector (Brandel, MD). Polysome sedimentation graphs were generated with Prism (v10.6; GraphPad).

### RNA-Seq Analysis of Polysome Fractions

Total RNA or pooled fractions from heavy polysomes were isolated using Trizol (ThermoFisher, 15596026) using standard procedures, except the precipitant in the aqueous fraction was isolated using an RNeasy column (Qiagen, 74134). mRNA was isolated from these total fractions using polyA enrichment (Dynabeads mRNA DIRECT Micro Purification Kit, ThermoFisher 61021) according to the manufacturer’s protocol, then library preparation was performed with Ion Total RNA-Seq Kit v2.0 (Thermo Fisher). Library size, concentration, and quality was assessed using a 2200 TapeStation (Agilent). Libraries were sequenced on Proton sequencer (ThermoFisher Scientific) with a PI chip and the Ion PI Hi-Q Sequencing 200 Kit for 520 flows. Ion Torrent reads were processed using combined Human Hg19 and KSHV (Accession GQ994935) reference transcriptomes. The KSHV genome was manually re-annotated with the transcript definitions from KSHV2.0 ^2^ reference transcriptome and the Quasi-Mapping software Salmon ^48^. Normalized <u>c</u>ounts <u>p</u>er <u>m</u>illion (cpm) were estimated for individual transcripts using the R package limma ^49^. Two biological replicates were combined as a geometric mean ^50^. The transcripts were ordered by abundance and the mean, standard deviation (SD), and Z-score were calculated using a sliding window of 200 transcripts of similar abundance ^50,51^. The most abundant 100 and the least abundant 100 transcripts used the mean and SD of the adjacent bin. Translational efficiency (TE) of a transcript treatment was determined by the formula TE = log_2_(polysome/total). The change in translational efficiency (DTE) = TE_Torin_-TE_DMSO_. We scored the difference in translational efficiencies by using a sliding-window to calculate a Z-score of each detected transcript compared to the surrounding 200 transcripts of similar abundance as measured by count per million (CPM).

### Polysome RT-qPCR

iSLK.219 were reactivated with 1 µg/mL doxycycline for 48 h. Torin or DMSO was added 2 h prior to harvest in high salt lysis buffer as described above. Select fractions were mixed 1:1 with Trizol and isolated as per manufacturer’s directions except that 30-60 µg of GlycoBlue Co-Precipitant (Invitrogen, AM9516) and 100 ng of *in vitro* transcribed luciferase DNA (T7 HiScribe, NEB E2040S) was added to the aqueous fractions during isopropanol precipitation ^47^. The resulting pellet was resuspended in water and reverse transcribed using Maxima H minus reverse transcriptase with random primers. mRNA was normalized to luciferase spike to control for recovery. The quantity of mRNA detected in each fraction was then calculated as a percentage of the total detected in all fractions. The RNA recovery was controlled by subtracting the C_T_ of the luciferase spike, which was assumed to be constant, from the target C_T_. This ΔC_T_ value for each fraction was then subtracted from the ΔC_T_ of the top fraction of the gradient to determine the ΔΔC_T_. And transcript abundance (Q) was then calculated (Q=2^ΔΔCT^) for each fraction, and all fractions were summed. The total quantity of a transcript is represented as a proportion of the total amount of detected transcript as per ^47^.

### LC-MS/MS proteomics

iSLK.219 were transfected with siRNA as described above and seeded onto 6-well plates. For lytic samples, the following day cells were treated with 1 µg/mL doxycycline; latent cells were untreated. For Torin treatment, drug or DMSO control was spiked into the cells 24 h after addition of doxycycline. For all experiments the cells were washed once with ice-cold PBS, scraped, pelleted at 500 x *g* for 5 min, then the washed once more in ice-cold PBS. Afterwards the cell pellet was snap-frozen in liquid nitrogen and stored at -80^°^C until processing.

Protein was extracted from cell pellets using 150μL of a solution of 20mM TrisCl pH 7.5, 150mM KCl, 5mM MgCl2, 0.5% (v/v) NP□40, 0.5% (w/v) sodium deoxycholate, and 0.5X cOmplete protease inhibitor (Roche, 04693116001). To support degradation of DNA/RNA, 10μL of a nuclease solution (1U/μL benzonase) was added to the solution and incubated at 24°C for 10-minutes. After nuclease digestion, 50μL of a solution of 600mM TrisCl pH 7.5, 8% (w/v) SDS, and 10mM dithiothreitol (DTT) was added to the tissue lysate and incubated at 95°C for 5-minutes in a heating block. After cooling to 24°C, lysates were alkylated using 40mM chloroacetamide (final concentration) and subsequently quenched with DTT. Protein concentration of the resulting lysate was measured using a BCA assay (ThermoFisher Scientific). Protein cleanup prior to digestion was carried out using an adapted version of the previously described SP3 protocol ^52,53^. Specifically, 1μL of a mixture of Sera-Mag carboxylate SpeedBeads (CAT#45152105050250 and CAT#65152105050250, prepared as 100mg/mL stock solution in water) was added to 50μg of protein for each sample prior to 4X volumes of acetone. Mixtures were incubated on a Thermomixer at 37°C for 5-minutes at 800 RPM prior to centrifugation at 5,000 x g for 5-minutes. After spinning, the supernatant was discarded and beads rinsed using 800μL of an 80% (v/v) ethanol solution with pipetting to resuspend the beads. Mixtures were centrifuged at 5,000 x g for 5-minutes and the supernatant discarded prior to the addition of 1μg of trypsin/rLysC mix (Promega) in 100 μL of digestion solution (100 mM ammonium bicarbonate, 1mM CaCl2) and incubation at 37°C for 16-hours in a Thermomixer with mixing at 800 RPM. After digestion, peptides were centrifuged at 12,000 x g for 2-minutes and the supernatant recovered to a fresh tube containing 5μL of a 10% (v/v) solution of trifluoroacetic acid. To desalt peptides prior to mass spectrometry (MS) analysis, an HPLC system (Agilent 1290 Infinity II with a DAD module) equipped with a reversed-phase column (CORTECS T3 1.6μm, 2.1×50mm, Waters) was utilized. Specifically, peptides were injected to a column equilibrated at 4% mobile phase B (0.1% formic acid in acetonitrile), rinsed for 1.5-minutes at a flow rate of 1mL/min, and subsequently eluted for 0.8-minutes at 80% mobile phase B (mobile phase A = 0.1% formic acid in water). Desalted peptides were dried in a SpeedVac centrifuge and reconstituted in 1% (v/v) formic acid in water at a concentration of 1 μg/μL based on the 214nm UV signal from the HPLC desalting step.

For total proteome analysis of si-eIF3d treated cells, peptide samples were analyzed using a data-independent acquisition (DIA) acquisition routine on an Orbitrap Fusion Lumos mass spectrometer (MS) (ThermoFisher Scientific). Samples were introduced to the MS using an Dionex UltiMate liquid chromatography (LC) instrument (ThermoFisher Scientific) equipped with a trapping-analytical column setup. For injection, peptides were initially trapped using 95% mobile phase A (0.1% formic acid) on a 100 μm inner diameter x 3cm length column packed in-house with 1.9 μm Reprosil Pur C18 beads (Dr. Maisch, r119.aq.). Gradient elution of peptides was performed using a ramp of mobile phase B (80% v/v acetonitrile in 0.1% formic acid) on a 100 μm inner diameter x 25 cm length analytical column packed with 1.9μm Reprosil Pur C18 beads. For each injection, 1 μg of peptides were separated with an 80 minute LC linear gradient from 5% to 11% mobile phase B in 0.5-minutes, and to 34% B in 72.5-minutes at 400 nL/min, followed by rinsing and re-equilibration over the remaining 7-minutes. The analytical column outlet was coupled to a 20 μm inner diameter LOTUS electrospray tip (Fossil Ion Tech.). Each peptide sample was injected three individual times (80-minute acquisition time for each), with each acquisition covering analysis of a separate mass range (injection 1 = 430 - 550, 2 = 550 - 690, 3 = 690 – 930 m/z). Specifically, the Orbitrap Lumos MS was globally set to use a positive ion spray voltage of 2200 V, an ion transfer tube temperature of 275°C, a default charge state of 3, and an RF Lens setting of 45%. The complete duty cycle of the acquisition method consisted of two MS1 scans and two sets of windowed MS2 scans (order MS1 - MS2 - MS1 - MS2). For the first, low mass range injection, the initial MS1 scan covered a mass range of 425-555m/z at a resolution of 60,000 with an AGC target of 4e5 (100%) and a max injection time set to ‘Auto’. The following set of DIA MS2 scans covered a precursor range of 430-550m/z with an isolation window size of 4 m/z (0m/z overlap) for a total of 30 scan events. Each scan used an HCD energy of 30% and covered a defined mass range of 200-1800 m/z at a resolution of 30,000 with a normalized AGC target of 1000%, and the maximum injection time set to ‘Auto’. Loop control was set to ‘N’ with a value of 30 spectra. The next MS1 scan and following DIA MS2 used the same settings as the previous with the exception that the MS2 precursor mass range was set to 428-552m/z to give a 2 m/z stagger with the previous scan windows. For the second, medium mass range injection, the scan settings were the same as for the low mass injection with the following changes: 1. MS1 scan range of 545-695 m/z; 2. DIA-MS2 precursor scan ranges of 550-690m/z and 548-692 m/z; 3. Loop control number of spectra = 35. For the third, high mass range injection, the scan settings were the same as the others with the following changes: 1. MS1 scan range of 685-935 m/z; 2. DIA-MS2 precursor scan ranges of 690-930m/z and 686-934m/z; 3. DIA-MS2 isolation window of 8 m/z; 4. Loop control number of spectra = 30. All scan data were acquired in centroid mode. DIA-MS raw data were de-multiplexed using msconvert (ProteoWizard, peakPicking = ‘vendor’, demultiplex = overlap only, 10ppm, SIM as spectra) and processed using DIA-NN (version 1.8.2 beta 8) ^54^. Specifically, a representative human proteome fasta database (version 01/2024, 20,521 entries, includes non-human contaminants and the KHSV proteome) was provided to DIA-NN along with all of the raw data in order to perform an initial spectral library generation step (default settings for library generation, --min-pr-mz 430 --max-pr-mz 930 --min-pr-charge 2 --max-pr-charge 4, MBR disabled). The entire set of raw files for each mass range were then searched against the spectral library using the derived mass error and window settings with MBR enabled. Resulting report files were passed to the iq package (Lib.Q.Value = 0.01, Lib.PG.Q.Value = 0.01, Q.Value = 0.01, PG.Q.Value = 0.05) in R to generate estimates of protein abundance ^55^.

For proteome analysis of si-eIF3d samples treated with Torin or DMSO, peptide samples were analyzed as above using a DIA acquisition routine on an Orbitrap Fusion Lumos MS with minor modifications. Specifically, peptides were initially trapped on a pre-column (ACQUITY UPLC M-Class Symmetry C18 Trap, 180µm x 2cm, 5µm beads, Waters) and then separated by an analytical column (nanoEase M/Z Peptide BEH C18, 75µm x 25cm, 1.7µm beads, Waters). Samples were initially trapped for 3-minutes at a flow rate of 20µL/min of 1% mobile phase B (0.1% formic acid in acetonitrile). Gradient separation of trapped peptides started at an initial condition of 1% mobile phase B, ramped to 8% B in 1-minute, to 30% B in 45-minutes, to 80% B in 0.5-minutes, hold at 80% B for 2-minutes, ramp to 1% B in 0.5-minutes, and a final hold at 3% B for 11-minutes for a total run time of 60-minutes (mobile phase A = 0.1% formic acid in water, flow rate = 300nL/min). Each peptide sample was injected once covering analysis of the 430 – 550m/z mass range. The Orbitrap Lumos MS was globally set to use a positive ion spray voltage of 2000 V, an ion transfer tube temperature of 275°C, a default charge state of 3, and an RF Lens setting of 45%. The complete duty cycle of the acquisition method consisted of two MS1 scans and two sets of windowed MS2 scans (order MS1 - MS2 - MS1 – MS2). The initial MS1 scan covered a mass range of 415-565m/z at a resolution of 30,000 with an AGC target of 4e5 (100%) and a max injection time set to ‘Custom’ (54ms). The following set of DIA MS2 scans covered a precursor range of 430-550m/z with an isolation window size of 4m/z (0m/z overlap) for a total of 30 scan events. Each scan used an HCD energy of 30% and covered a defined mass range of 200-1800m/z at a resolution of 30,000 with a normalized AGC target of 1000%, and the maximum injection time set to ‘Custom’ (54ms). Loop control was set to ‘N’ with a value of 30 spectra. The next MS1 scan and following DIA MS2 used the same settings as the previous with the exception that the MS2 precursor mass range was set to 428-552m/z to give a 2m/z stagger with the previous scan windows. All scan data were acquired in centroid mode. Analysis of acquired MS data was performed as above with DIA-NN (version 2.1.0) and an updated human proteome database (version 05/2025, 20,514 entries, includes non-human contaminants and the KHSV proteome). Differential protein expression was determined using the R package DEqMS^56^. Volcano plots were then generated in Prism (v10.6; GraphPad).

### GSEA method

The results from DEqMS were used to generate ranked gene lists using the “t” statistic from lowest to highest. These ranked lists were analysed in GSEA 4.3.3 ^25^ using the Reactome canonical pathways in the Human Molecular Signatures Database (cp.c2.reactome.v2023.2.Hs.symbols) where we manually appended an additional “KSHV” gene set of as used in our proteomic analysis. We used a 10% FDR as a cut off for the gene sets presented in **Fig. 1E** and **Fig. 4B** ^25^. Bubble plots were generated with ggplot2 in R.

### Statistics

RT-qPCR, cholesterol quantification, and titering values were calculated in Excel (Microsoft). Values were imported into Prism (v10.6; GraphPad) for statistical analysis and graphing. *P* values are represented in the figures (*, *P* < 0.05; **, *P* < 0.01; ***, *P* < 0.001; ns, nonsignificant).

## Supporting information

Supplemental raw unprocessed images for western blots

## Data availability

The mass spectrometry proteomics data have been deposited to the ProteomeXchange Consortium via the PRIDE ^57^ partner repository with the dataset identifier PXD069250

## ACKNOWLEDGEMENTS

We are grateful for the support of Gerard Gaspard, Facility Manager of the Dalhousie University Cellular and Molecular Digital Imaging Core Facility for invaluable support in training, troubleshooting, and data analysis. We are grateful to Don Ganem (University of California San Francisco) for the gifts of KSHV LANA antiserum, iSLK cells, and iSLK.219 cells. We thank Janani Krishnan (Dalhousie University) for technical support and Barbara Karten (Dalhousie University) for helpful advice. This work was supported by Canadian Institutes for Health Research Project Grant (PJT-173510) to C.M. The funders had no role in study design, data collection, and interpretation, or the decision to submit the work for publication.

## SUPPORTING INFORMATION

**S1 Table** – Source data related to Fig. 1C

**S2 Table** – Source data related to Fig. 1D

**S3 Table** – Source data related to Fig. 3A

**S4 Table** – Source data related to Fig. 3B

**S5 Table** – Source data related to Fig. 3C

**S6 Table** – Source data related to Fig. 4A

**S7 Table** – Source data related to Fig. 5A

**S8 Table** – Source data related to Fig. 5B

**S9 Table** – Source data related to Fig. 5C

**S10 Table** – Source data related to Fig. 5D

**S11 Table** – Source data related to Fig. 6A

**S12 Table** – Source data related to Fig. 6B

**S13 Table** – Source data related to Fig. 1C

**S1_raw_images** – Raw blot images for westerns in Fig 1A, 2A, 2B, and 8B.

