## Supplemental raw unprocessed images for western blots for "eIF3d drives cholesterol biosynthesis required for KSHV lytic replication"

### Fig 1A - Original blot images

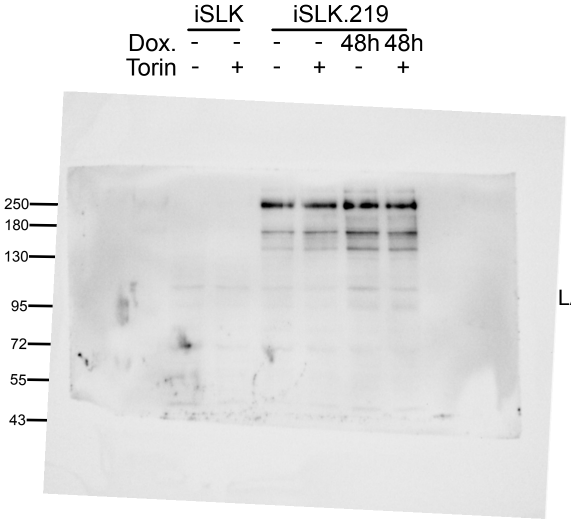

LANA

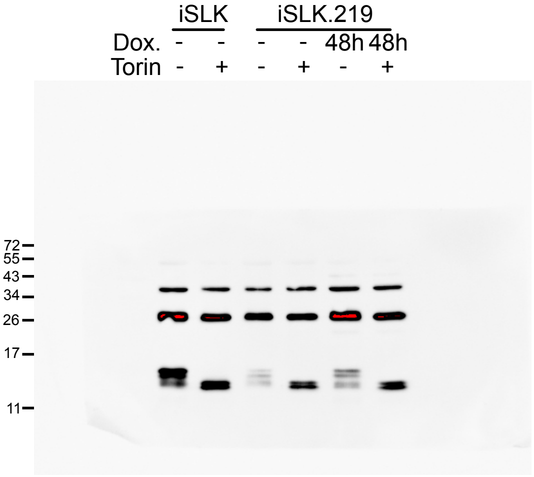

β-actin
Reprobe on S6
Reprobed for 4E-BP

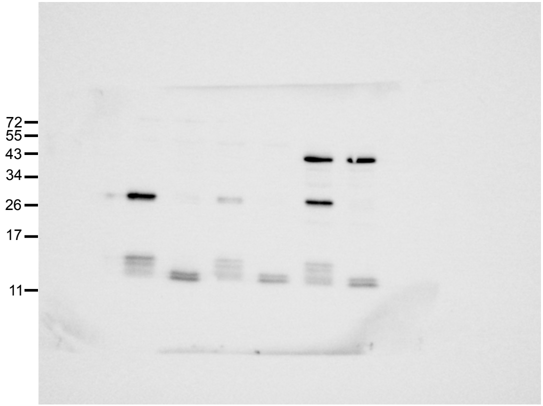

ORF57
Reprobe on p-S6

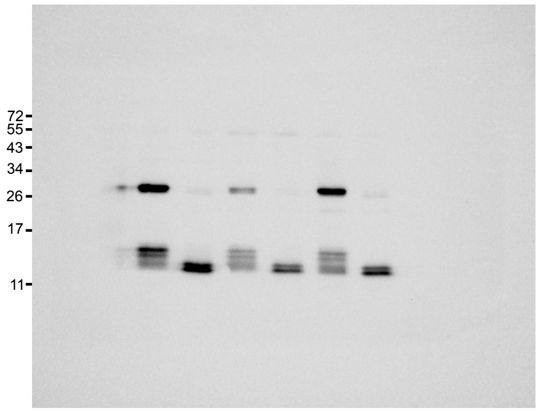

p-S6
Reprobed for 4E-BP

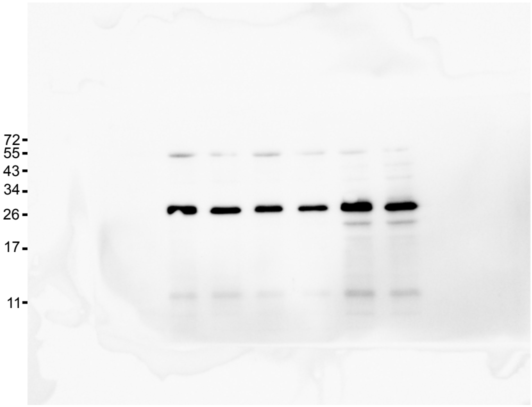

S6

### Fig 1A - Cropped

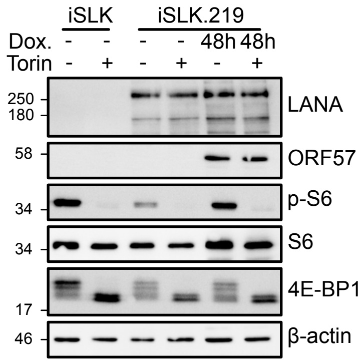

### Fig 2A - Original blot images

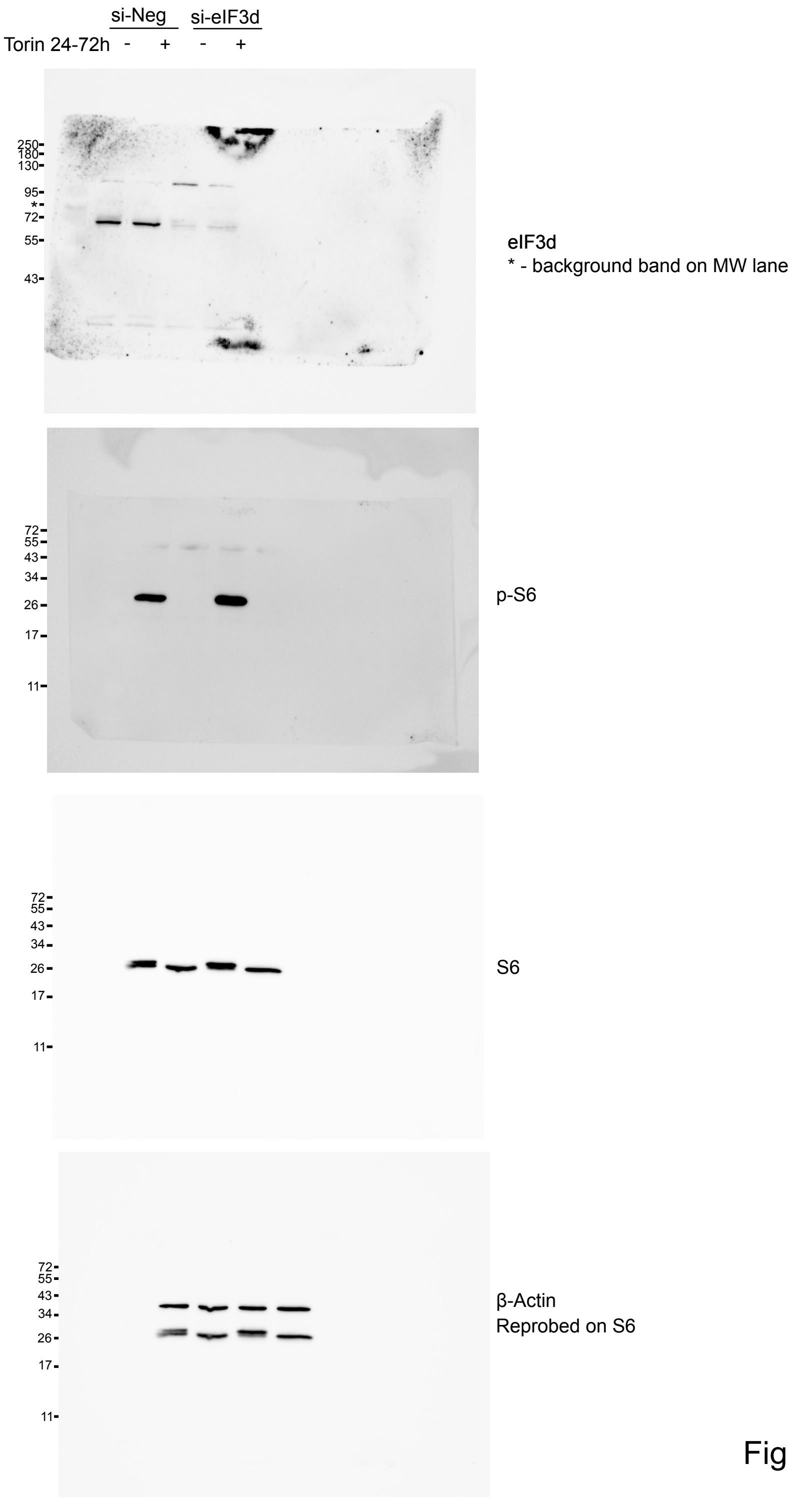


### Fig 2A - Cropped

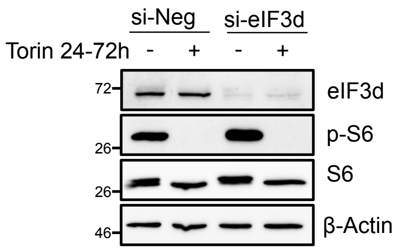

### Fig 2B - Original blot images

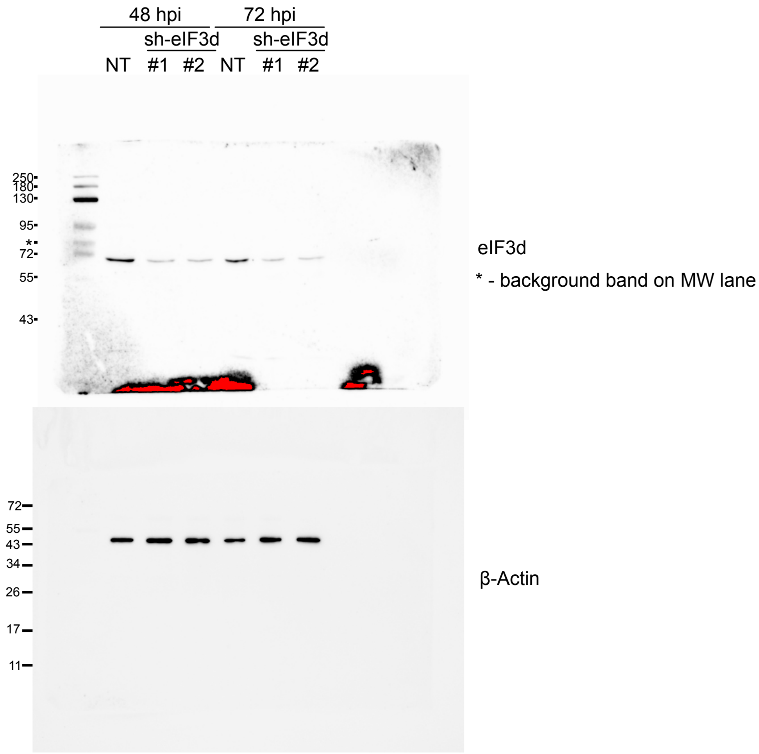


### Fig 2B - Cropped

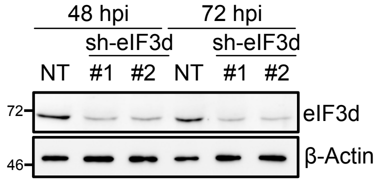

### Fig 8B - Original blot images

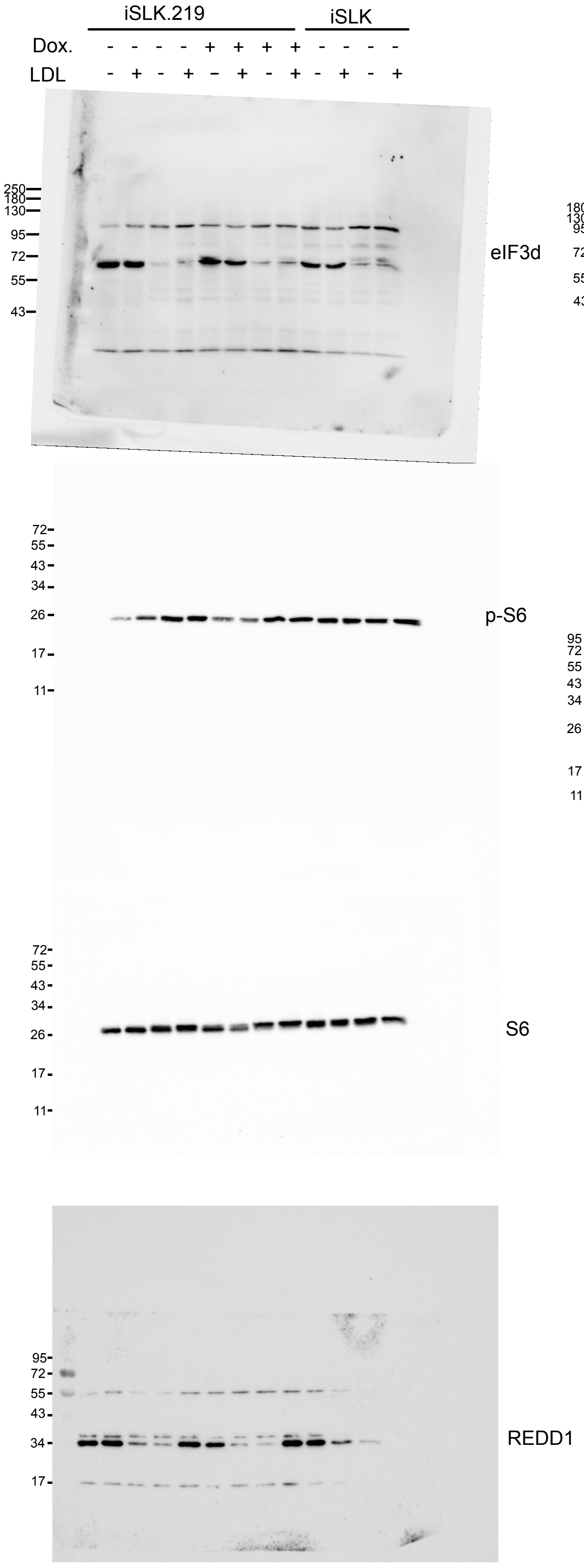


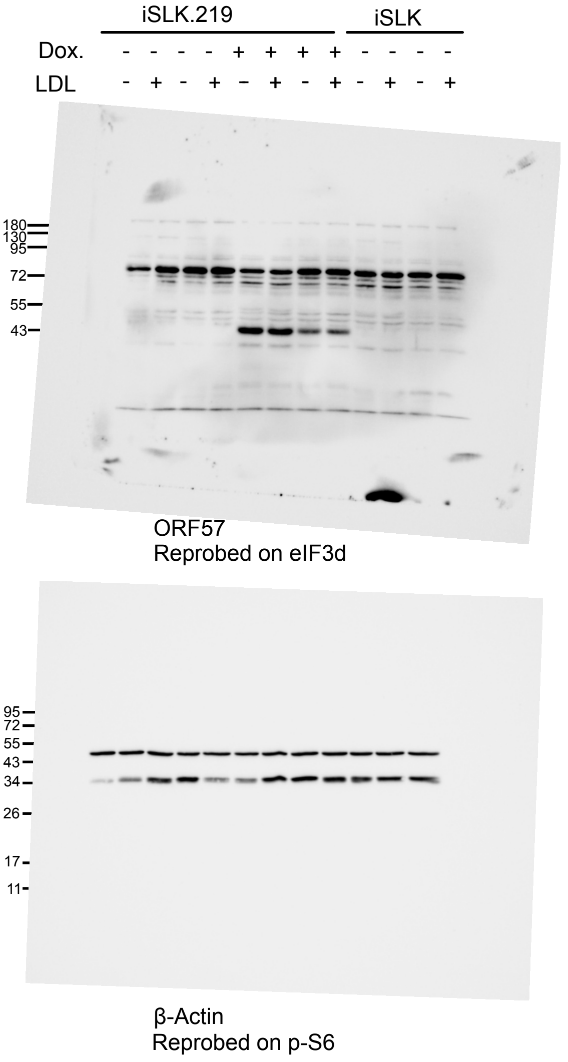


### Fig 8B - Cropped

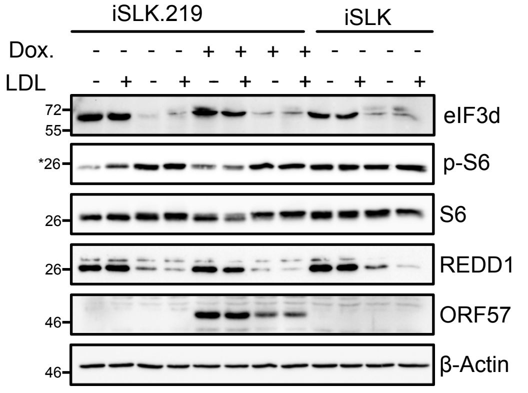
